# Ultrastructural analysis of wildtype and RIM1α knock-out active zones in a large cortical synapse

**DOI:** 10.1101/2022.03.04.482996

**Authors:** Katharina Lichter, Mila Marie Paul, Martin Pauli, Susanne Schoch, Philip Kollmannsberger, Christian Stigloher, Manfred Heckmann, Anna-Leena Sirén

## Abstract

Rab3A-interacting molecule (RIM) is crucial for fast Ca^2+^-triggered synaptic vesicle (SV) release in presynaptic active zones (AZ). While loss of RIM1α impairs long-term plasticity at hippocampal giant mossy fiber boutons (MFB), it remains unclear how AZ ultrastructure is altered. We investigated MFB AZ architecture in 3D using electron tomography of rapid cryo-immobilized acute brain slices in RIM1α^-/-^ and wild-type mice. In RIM1α^-/-^, AZs are larger with increased synaptic cleft heights and with a three-fold reduced number of tightly docked SVs (0-2nm). The distance of tightly docked SVs to the AZ center is increased from 110 to 195 nm, and the width of their electron dense material between outer SV membrane and AZ membrane is reduced. Furthermore, the SV pool in RIM1α^-/-^ is more heterogeneous. Thus, RIM1α, beside its role in tight SV docking, is crucial for synaptic architecture and vesicle pool organization in MFBs.

## Introduction

Rab3A-interacting molecules (RIM) form evolutionarily conserved, presynaptic scaffold complexes localized at the active zone (AZ) mesoscale (Emperador-Melero and Kaeser, 2020; Goodsell et al., 2020). In synaptic contacts of vertebrates and invertebrates, RIM and homologues facilitate synaptic transmission and its potentiation for short or long-term information storage (Castillo et al., 2002; Kintscher et al., 2013; Müller et al., 2012; Paul et al., 2022; Schoch et al., 2002; Stigloher et al., 2011).

RIM contains 5 core protein domains (Zinc finger, PDZ, C_2_A, PxxP, C_2_B). Via the N-terminal Zinc finger it binds to Rab3A (Wang et al., 1997), Munc13-1 (Andrews-Zwilling et al., 2006; Deng et al., 2011) and via PxxP the scaffold protein RIM-BP (Hibino et al., 2002; Wang et al., 2002). Its PDZ-, C_2_A- and C_2_B-domain bind to P/Q- and N-type voltage gated calcium channels (VGCCs) (Deng et al., 2011; Han et al., 2011; Kaeser et al., 2012; Kaeser et al., 2011; Kiyonaka et al., 2007; Miki et al., 2007). Additionally, the C_2_B-domain interacts with the presynaptic membrane (de Jong et al., 2018) and liprin-α3 (Schoch et al., 2002).

RIM1α is the major RIM isoform in the hippocampal mossy fiber / CA3 region (Schoch et al., 2006). In electrophysiological studies, RIM1α deficiency (RIM1α^-/-^) impaired long-term potentiation (LTP) of giant mossy fiber boutons (MFB) CA3 pyramidal neuron synapses (Castillo et al., 2002). Plasticity in MFBs is expressed presynaptically (Nicoll and Schmitz, 2005; Vandael et al., 2020; Vyleta and Jonas, 2014). Within the tri-synaptic hippocampal circuit, giant MFB synapses function as conditional detonators to time and control downstream postsynaptic activity (Henze et al., 2002; Vyleta et al., 2016) and are, thus, indispensable for memory. Furthermore, giant MFB synapses are characterized by a large pool of release-ready SVs, low release probability and loose coupling distance between VGCC and Ca^2+^ sensor (Jonas et al., 1993; Vyleta and Jonas, 2014).

Like the physiological properties, the morphology of giant MFBs is diverse (Rollenhagen et al., 2007; Wilke et al., 2013). Recently, a function to structure relationship was demonstrated for the association of post-tetanic potentiation (PTP) (Vandael et al., 2020), with an increase of the readily releasable pool (RRP) and quantified via “flash-and-freeze” electron microscopy (EM) (Borges-Merjane et al., 2020; Vandael et al., 2020; Watanabe et al., 2013). On an ultrastructural level, EM investigations of RIM1α deficiency showed a reduction of the docked SV pool and SV tethering in rapid-cryoimmobilized synaptic preparations of mammalian neurons (Fernandez-Busnadiego et al., 2013).

To further clarify the molecular architecture of the giant MFBs and their complex SV pools, we studied AZs in RIM1α^-/-^ mice. We used systematic, highly standardized near to native electron tomography for 3D nanoscopic quantification of synapses in rapid cryo-immobilized acute hippocampal slices of male adult RIM1α^-/-^ mice and age matched wildtype littermates. We answer the following questions:

1. Do the size of giant MFB AZs and the synaptic cleft width depend on RIM1α?
2. Are number and position of docked SVs influenced by RIM1α and is the electron dense material below the docked SVs altered?
3. Is the organization of the SV pool up to 200 nm from the presynaptic membrane changed in RIM1α^-/-^?

## Results

### Targeted cutting for imaging of giant MFB AZs

Giant MFB AZs are highly heterogenous within the mossy fiber (MF) tract and along the dorsoventral hippocampal axis (Kheirbek et al., 2013; Pauli et al., 2021). Thus, we used a standardized protocol for nanoscopic AZ analysis of brains of three male RIM1α^-/-^ and three wildtype littermates to prepare acute slices of the left dorsal hippocampus (Figure 1A, B). To avoid fixative induced alterations of AZ architecture (Korogod et al., 2015; Weimer, 2006; Zhao et al., 2012) rapid cryo-immobilization and freeze substitution were used (Figure 1C). We applied targeted ultramicrotomy to clearly identify supra-pyramidal giant hippocampal MFB to CA3b spine head synapses (Figure 1D). Electron microscopic (EM) tomography was carried out on 250 nm semi-thin hippocampal resin sections in high resolution and magnification to obtain a 3D morphological dataset for analysis of MFB AZs (Figure 1E, F). All EM tomograms were reconstructed with a fiducial free patch tracking protocol and annotated as individual IMOD models (Suppl. Figure 1, N = 30 AZs in wildtype, N = 32 AZs in RIM1α^-/-^).

**Figure 1.**
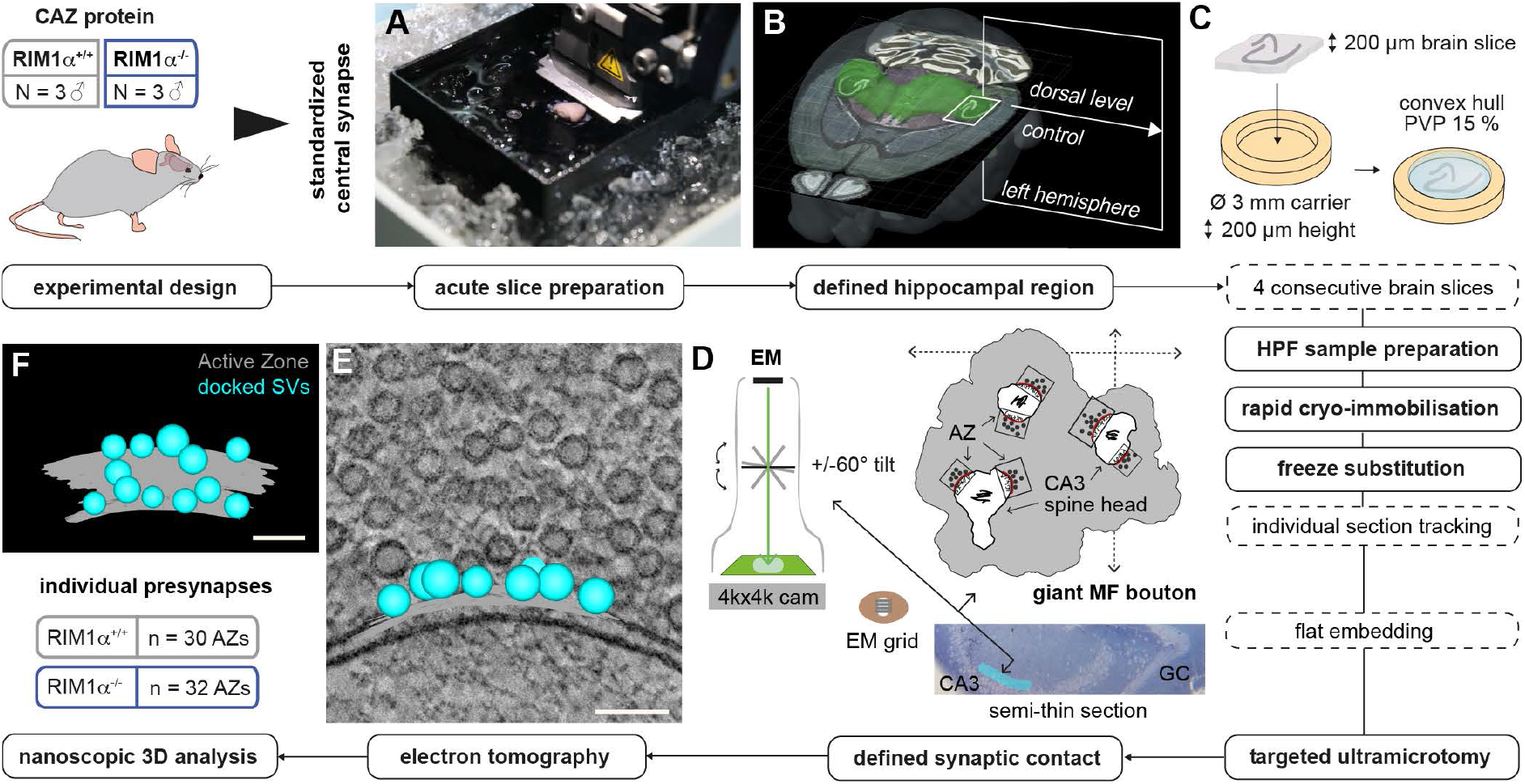
Schematic workflow for standardized ultrastructural analysis of presynapses in giant hippocampal MF boutons. **A** Young adult wildtype (RIM1α^+/+^, grey, N = 3) and RIM1α knock-out mice (RIM1α^-/-^, blue, N = 3) were used for systematic analysis of three-dimensional AZ architecture in standardized central synapses to decipher morphological phenotypes of cytomatrix AZ proteins (CAZ). After decapitation, brains were removed immediately for acute slicing in ACSF using a vibratome. **B** Horizontal section from Allen Mouse Brain Atlas illustrates the region of interest in the dorsal left hippocampus from which at least 4 consecutive 200 µm thick slices were obtained. **C** Slices were transferred in high-pressure freezing (HPF) carriers of 3 mm diameter and 200 µm height (Type A) filled with PVP 15% as cryoprotectans. **D** Summary of sample processing including rapid cryo-immobilisation, freeze substitution and targeted ultramicrotomy for electron tomography of defined synaptic contacts of giant hippocampal MFBs. **E** Reconstructed electron tomographic subvolume of a MF synaptic contact. Presynaptic active zone (AZ, grey) and docked synaptic vesicles (SVs, cyan) are highlighted. **F** Nanoscopic 3D analysis of individual MF presynapses was performed using ETomo/ IMOD and Python software packages in a total number of 62 synaptic contacts in RIM1α^+/+^ and RIM1α^-/-^ animals. Scale bars: (E, F) 100 nm.

### AZ size and synaptic cleft width are increased in RIM1α^-/-^

We identified morphological characteristics of giant MFBs such as dense filling of boutons with SVs including dense-core and large clear vesicles (Rollenhagen et al., 2007)(Figure 2A-D). 2D synaptic profiles with clear and intact pre- and postsynaptic membranes were used for image acquisition of tilt series. Synaptic contacts were identified as presynaptic membranes with corresponding postsynaptic filaments (PSF) at the postsynaptic membrane of CA3b spine heads (Figure 2E-H). In contrast to the postsynaptic density of classically fixed electron microscopic samples, single electron dense, partially low-contrast filamentous structures could be differentiated (Figure 2E1-H1). RIM1α^-/-^ giant MFB to CA3b spine head AZs were significantly larger than wildtype AZs (Figure 2I-J, see Table S1 for all numerical and statistical values not stated in the text). Furthermore, the distribution of 2D AZ profile lengths was more variable in RIM1α^-/-^ (Figure 2K). Next, we determined the maximum extent of individual AZ areas in 3D which was also significantly increased in RIM1α^-/-^ mice from 602.31 nm in wildtype to 802.50 nm in RIM1α^-/-^ (Figure 2L).

**Figure 2.**
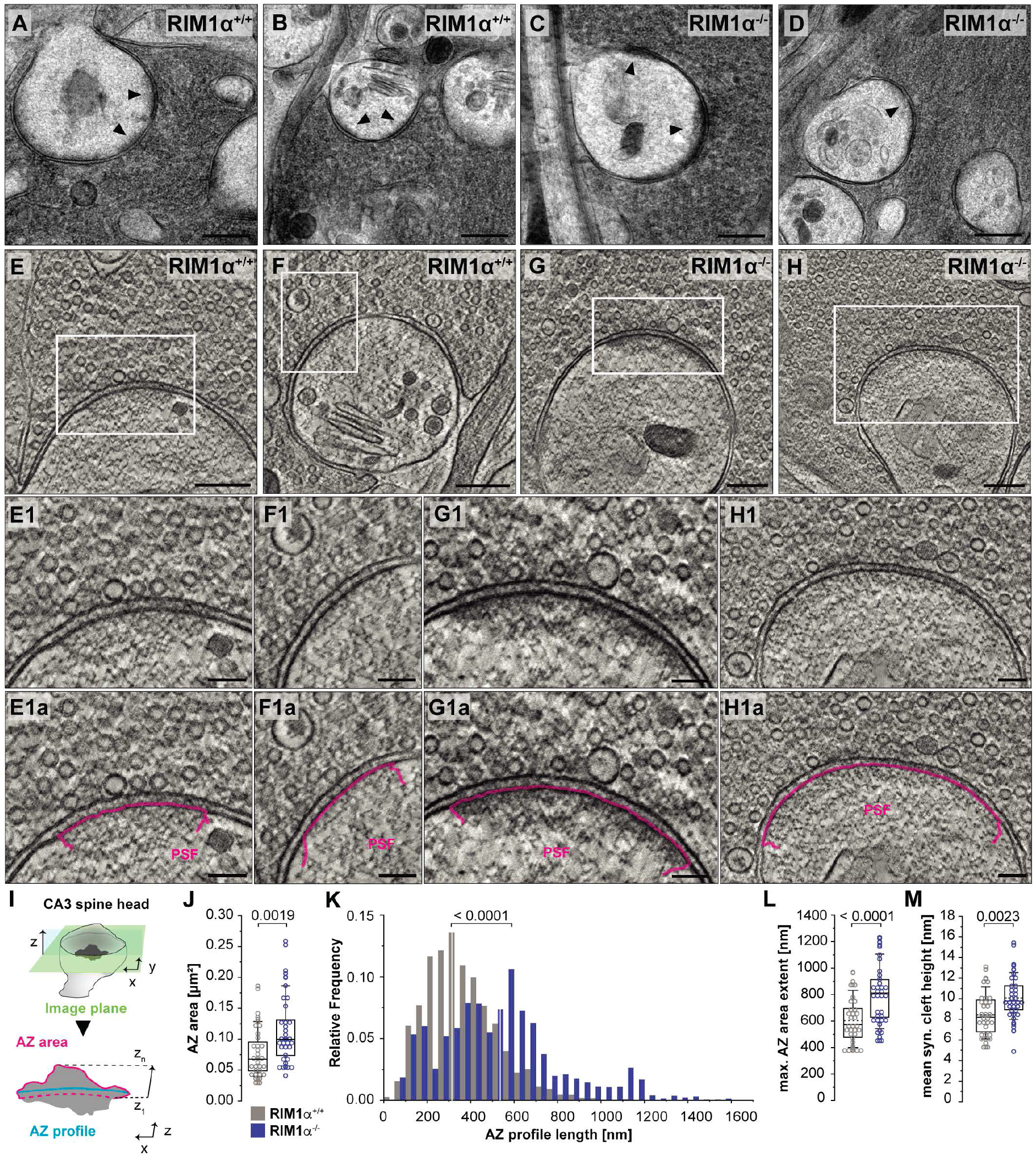
Ultrastructural 3D analysis reveals larger hippocampal MFB AZs in RIM1α^-/-^ mice. **A-D** Exemplary electron micrographs show overviews of large CA3 MFs within the left dorsal hippocampus of RIM1α^+/+^ (A, B) and RIM1α^-/-^ mice (C, D). Black arrow heads highlight individual presynaptic AZs used for electron tomography. **E-H** Selected electron tomographic slices of individual hippocampal MFBs (Figure 2 A-D) in RIM1α^+/+^ (E, F) and RIM1α^-/-^ (G, H). White boxes highlight regions enlarged in E1-H1. **E1-H1** Enlarged view indicated by white boxes in E-H show presynaptic AZs of hippocampal MFBs in RIM1α^+/+^ (E1, F1) and RIM1α^-/-^ mice (G1, H1). In E1a-H1a, postsynaptic filaments (PSF) are highlighted in magenta. **I** Illustration of selected image plane (green) for visualization of presynaptic MF AZs on CA3 spine heads. Measurements of AZ area and AZ profile are indicated below. **J** Summary graphs of AZ area in RIM1α^+/+^ (grey, n = 30 AZs, 3 animals) and RIM1α^-/-^ (blue, n = 32 AZs, 3 animals). Throughout this manuscript, horizontal lines in box plots represent median; boxes quartiles; whiskers 10^th^ and 90^th^ percentiles; scatter plots show individual data points for each group. P-values are indicated above summary graphs. **K** Histograms of AZ profile lengths show broader distribution in RIM1α^-/-^ (n = 12,750 AZ profile lengths) than RIM1α^+/+^ (n = 13,991 AZ profile lengths). **L, M** Summary graphs of the maximal AZ extent (L) obtained and synaptic cleft height (M) calculated based on IMOD models in both groups. Scale bars: (A-D) 250 nm, (E-H) 250 nm, (E1-H1) 100 nm.

We annotated the synaptic cleft volume below the defined presynaptic area with its full extent to the AZ edges in a separate 3D object and normalized it to the individual AZ area. The height of the synaptic cleft was increased in RIM1α^-/-^ compared to wildtype controls (Figure 2M). Furthermore, we observed that the synaptic cleft was slightly smaller laterally towards the AZ edges and depending on the curvature of the spine head membranes, but within the previously reported range (Savtchenko and Rusakov, 2007).

We observed more mitochondria near the AZ and the perisynaptic zone in RIM1α^-/-^ (Figure S2A, B). To quantify this, we used a 2D spatial stereology with a systematic grid lattice and equal magnification (Figure S2C). The total mitochondrial area within 100-500 nm of the AZ membrane and its perisynaptic zone was increased in RIM1α^-/-^ AZs compared to wildtype (Figure S2D).

### Reduction of the docked SV pool and diminution of the SV-attached electron dense material at RIM1α^-/-^ AZs

We individually measured all SVs within 0-10 nm to the AZ membrane in high magnification (Figure 3A, Videos SA1-SA4). 0-10 nm covers the range of the SNARE proteins. SVs of 0-2 nm distance to the presynaptic membrane with prominent electron dense material (EDM) below the SV were defined as ‘docked’ (Imig et al., 2014). In RIM1α^-/-^, docking was greatly reduced (Figure 3B). In both genotypes, we observed AZs without any docked SVs. Since previous studies differ in the definition of docked SVs (Borges-Merjane et al., 2020; Neher and Brose, 2018; Rothman et al., 2017; Tang et al., 2016), we quantified the number of docked SVs in 0-5 nm distance. Here, the number of docked SVs in RIM1α^-/-^ AZs was also reduced as compared to wildtype AZs (Figure 3B). As we found no difference of docked SV numbers within 2.1-5 nm distance (Figure 3B), we attribute this difference to the 0-2 nm fraction. Furthermore, SV numbers within 5-10 nm distance from the membrane, where SVs are tethered to the membrane via EDMs (Fernandez-Busnadiego et al., 2013), were unchanged (Figure 3B). Interestingly, SVs within the 0-10 nm zone were nearly 2-fold further from the AZ membrane in RIM1α^-/-^ than in wildtype.

**Figure 3.**
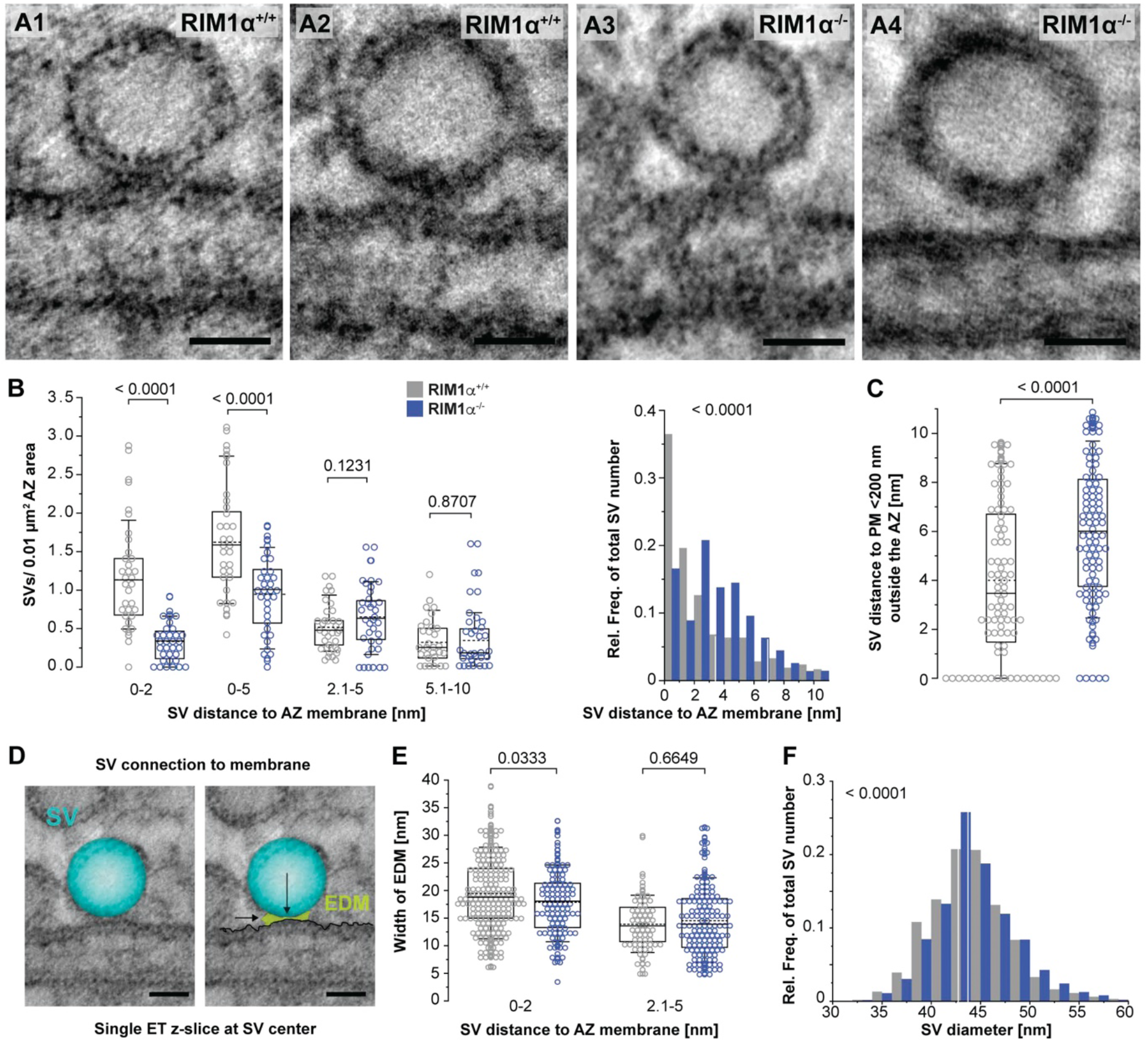
Altered docking of synaptic vesicles at RIM1α^-/-^ giant MFB AZs. **A** Electron tomographic slices of individual SVs depicted at their centers at the presynaptic membrane in RIM1α^+/+^ (A1-A2) and RIM1α^-/-^ (A3-A4). **B** Summary graphs of normalized SV numbers within 0-2 nm, 0-5 nm, 2.1-5 nm and 5.1-10 nm distance to the AZ membrane in RIM1α^+/+^ (grey) and RIM1α^-/-^ (blue) and histograms of the distance distribution of SVs and the AZ membrane within 0-10 nm. **C** Summary graph of outer docked and tethered SVs (i.e. < 200 nm to AZ edges) within 0-10 nm distance to the AZ membrane in both groups. **D** Visualization of a 3D docked SV (cyan) at its center with electron dense material (EDM, green) connecting SV and the presynaptic membrane in an electron tomographic slice in front view. **E** The maximal EDM width in docked SVs of 0-2 nm distance decreases in RIM1α^- /-^ AZs compared to controls. SVs of 2.1-5 nm distance, connected via EDM to the AZ membrane, don’t differ in their maximal EDM width. **F** SVs in the total SV pool (0-200 nm) have an increased diameter in RIM1α^-/-^ AZs compared to controls. Medians are indicated as white lines in histogram bins throughout the manuscript. Scale bars: (A1-A4, D) 25 nm.

In the nematode *Caenorhabditis elegans*, presynaptic dense projections localize SVs via an interaction of the RIM homologue *unc-10* at the neuromuscular junction (NMJ) which may include tethered and docked SVs in a perisynaptic zone (Stigloher et al., 2011). We quantified any SV below 10 nm to the membrane and within 200 nm perisynaptic distance to the AZ (Figure 3C). In both genotypes, we rarely detected SVs in a tethered or docked status outside of the AZ. The median SV distance to the membrane was significantly higher in RIM1α^-/-^ (Figure 3C) which may indicate a compromised ectopic release machinery in the absence of RIM1α.

Due to the distinct distribution of docked SVs between 0-2 nm in both genotypes, we measured the EDM connecting SV and membrane (Harlow et al., 2001). To our current knowledge, there has not been any systematic quantification of rapid cryo-immobilized EDM in docked SVs of 0-5 nm distance in central mammalian synapses (of acute brain slices) so far. We measured the EDM width at the SV center between 0-5 nm at high-resolution and normalized it to the corresponding SV diameter. Although our near-to-native tissue preparation and 3D tomogram reconstruction facilitated detection of thin electron dense filaments (Figure 3, Videos SA1-SA4), annotation of individual filaments remains highly subjective. Thus, we focused on the longest continuous part of the EDM (Figure 3D). In RIM1α^-/-^ AZs, the EDM below docked SVs in 0-2 nm is smaller compared to wildtype AZs (Figure 3E). EDM widths of SVs connected to the membrane in 2.1-5 nm distance did not differ (Figure 3E), however, the quantified EDM was significantly smaller in both genotypes compared to SVs in 0-2 nm distance (Figure S3, p < 0.0001). Normalized calculations of the EDM to its corresponding SV diameter further supported these findings (Figure S3). The overall diameter of SVs within 0-200 nm in RIM1α^-/-^ was larger compared to wildtype SVs (cut-off for radius: 30 nm, Figure 3F).

### Delocalization of the docked SV pool in RIM1α^-/-^

We systematically profiled individual docked SVs upon their five nearest neighboring docked SVs, their position to both AZ edges in the EM tomogram, and their distance to the 3D center of mass (c.o.m.) of the AZ (Figure 4A) (Butola et al., 2021; Kusick et al., 2020). In RIM1α^-/-^AZs, the distance of the index docked SV to its five nearest neighboring docked SVs is largely increased compared to controls (Figure 4B, Table S1). Using the 2D AZ profile length as a reference of the docked SV position (Figure 4A, (Kusick et al., 2020), the distance of the projected SV center onto the AZ profile to both corresponding edges was larger in RIM1α^-/-^(Figure 4C, D). Due to the huge AZ area increase in RIM1α^-/-^, the calculations were further normalized to the corresponding AZ profile (Figure 4A) and the ratio of both distances was compared: Docked SVs in RIMα^-/-^AZs are localized nearer towards the AZ edges compared to controls (Figure 4E). Docked SVs in wildtype AZs are located closer to the AZ c.o.m. (Figure 4E). As giant MFB AZs are complex in 3D, the c.o.m. of the AZ is one possible standardization to report spatial differences (Mrestani et al., 2021). The distance of an individual docked SV to the 3D c.o.m of the AZ was increased nearly two-fold in RIM1α^-/-^compared to wild-type (Figure 4E).

**Figure 4.**
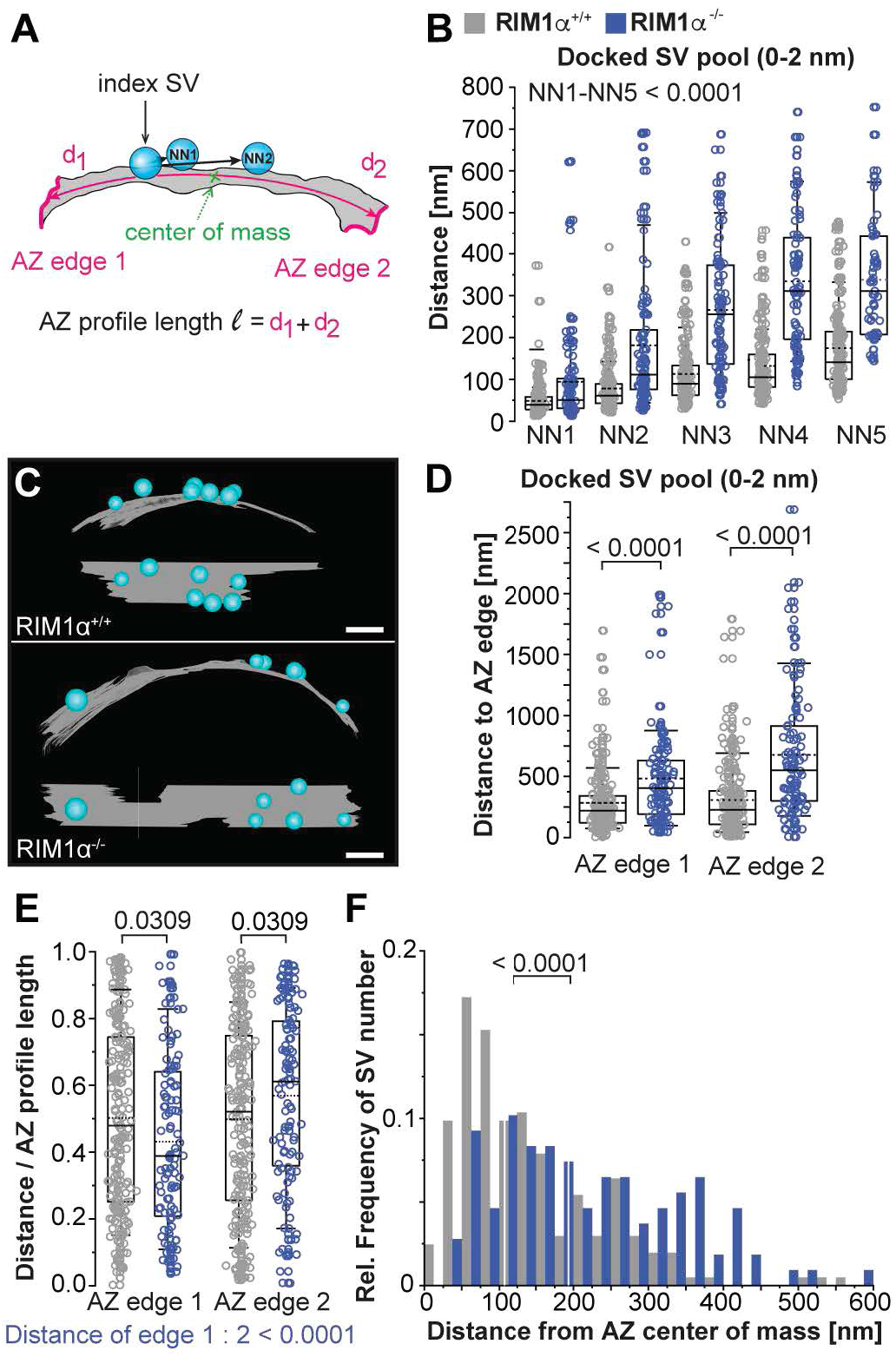
Delocalization of the docked SV pool (0-2 nm) at RIM1α^-/-^ giant MFB AZs. **A** Visualization of standardized distance measurements for the docked SV pool (0-2 nm distance to AZ membrane) at giant MFB AZs. The docked index SV (cyan) is localized to the 5 nearest neighboring docked SVs at the AZ membrane. Further, the position of the index SV within the AZ is computed via its distances (d_1_, d_2_) to corresponding AZ edges (magenta) and the 3D AZ center of mass (c.o.m). **B** Individual neighboring docked SVs are wider distributed at the AZ membranes in RIM1α^-/-^ AZs than in controls. **C** Exemplary IMOD models of the docked SV pool at the AZ membrane of RIM1α^- /-^ giant MFB AZs and controls. **D** Individual docked SVs show an increased absolute distance to corresponding AZ edges 1 and 2 compared to controls. **E** Normalized relative distances to the AZ profile length l at the SV center point indicate a distribution of docked SVs in RIM1α^-/-^ AZs which is shifted from the AZ center compared to controls. **F** The distance of docked SVs to the AZ c.o.m is increased and widely distributed in RIM1α^-/-^ AZs compared to controls.

### Heterogeneous distribution of SVs within the pool up to 200 nm in RIM1α^-/-^

The docked SV pool of giant MFB AZs is largely re-organized in the absence of RIM1α. Is this associated with a reduced SV pool? The total SV pool size (0-200 nm) positively correlated with the area of RIM1α^-/-^ and wildtype AZs (F = 1.699, p = 0.192, F-test; Figure 5A left panel). At 0-50 nm distance to the presynaptic membrane this correlation was still present, however, correlation strength was decreased in RIM1α^-/-^(Figure 5A, right panel). In RIM1α^-/-^MFB AZs, the total SV pool was reduced. Taken the median SV diameter of 43-44 nm at giant MFB AZs plus a zone of SV proteins on the outer SV membrane, it is reasonable to differentiate the SV pool in 2D distance fractions of 50 nanometers (Kaeser and Regehr, 2017). In RIM1α^-/-^giant MFB AZs, all four fractions showed a significant reduction in SV number per 0.01µm^2^ AZ area (Figure 5B). In both genotypes, most SVs were found within 101-150 nm distance to the presynaptic membrane (Figure 5C, Figure S5).

**Figure 5.**
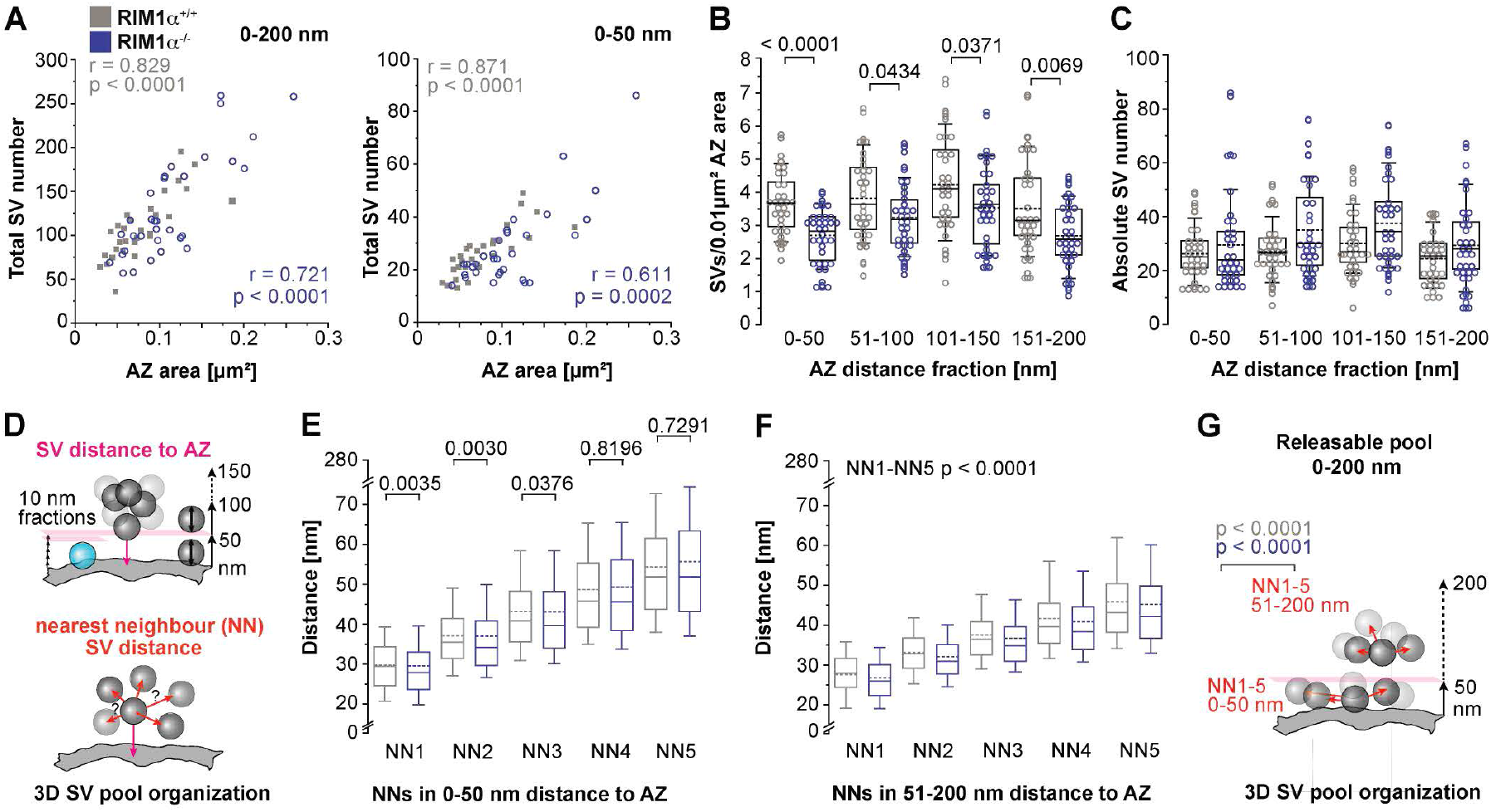
Distinct organization of the synaptic vesicle pool in RIM1α^-/-^ giant MFB synapses. **A** Spearman correlation coefficient of the total SV number in 0-200 nm (left panel) or 0-50 nm (right panel) distance to the presynaptic membrane and the area of individual AZs in RIM1α^+/+^ (grey) and RIM1α^-/-^ (blue) mice. **B, C** Summary graphs of normalized SV numbers (B) and absolute SV numbers (C) within 0-50 nm, 51-100 nm, 101-150 nm and 151-200 nm distance to the AZ membrane in both groups. **D** Illustration of 2D and 3D SV pool organization analyses in distance fractions of 10 or 50 nm distance from the AZ membrane (upper panel) and using nearest neighbor analysis (NN, lower panel). **E, F** Summary graphs of the distance from the AZ membrane for NN1-5 within 0-50 nm (E) and 51-200 nm (F) to the AZ. **G** Reserve pool - 3D illustration of the NN1-5 SV pool analysis within 0-50 nm and 51-200 nm distance to the AZ membrane.

We next performed the nearest neighbor (NN) analysis of surrounding SVs in 0-50 nm and 51-200 nm distance to the presynaptic AZ membrane (Figure 5D). In 0-50 nm distance to the AZ, NN 1-3 distances were smaller in the RIM1α^-/-^ SV pool (Figure 5E, Figure S5). In 51-200 nm even NN 1-5 distances were smaller (Figure 5F, Figure S5). Interestingly, the overall NN distance in in the more distant SV pool (51-200 nm) was significantly smaller than in the SV pool in 0-50 nm distance (for both genotypes: p < 0.0001, Mann-Whitney Rank Sum Test, Figure 5G).

## Discussion

The present study introduces the first quantitative 3D analysis of nanoscopic AZ architecture in giant MFBs using electron tomography of rapid cryo-immobilized acute brain slices. Since ablation of RIM1α is sufficient to abolish LTP in MFBs (Castillo et al., 2002), we concentrated on AZs in MFB to CA3b spine head synapses of adult male RIM1α knockout and wildtype littermates instead of RIM1/2 knockout mice (Kaeser et al., 2011). From a general perspective, our wildtype data of MFB AZs in acute brain slices such as AZ length, SV diameter, number of tightly docked SVs and the percent occupancy of AZ area with mitochondria fit with recent electron tomographic data from hippocampal slice cultures (Maus et al., 2020). RIM1α^-/-^ AZs showed an expansion of area and synaptic cleft, reduced docked SV pool and altered SV docking apparatus, lateralization of docked SVs towards the AZ edge, and a distinct SV pool organization up to 200 nm distance to the presynaptic membrane. These findings imply that RIM1α is essential for AZ nanoarchitecture and SV arrangement in giant MFBs.

### AZ size and synaptic cleft width depend on RIM1α

We found a striking increase of AZ size, 2D profile length and maximum extent in the MFB to CA3 spine head synapses in RIM1α^-/-^. These data are in accordance with electron tomography of cryo-immobilized cortico-cerebral synaptosomal preparations (Fernandez-Busnadiego et al., 2013). By targeted cutting and focusing on one defined synapse type, we obtained highly standardized data. Nevertheless, the AZ dimensions showed remarkable variability in wildtype which was further augmented in RIM1α^-/-^. On average AZ area increased by 30% in RIM1α^-/-^. Furthermore, the shape of an individual AZ showed a high complexity (Figure S1) which is in line with other studies (Pauli et al., 2021; Zhao et al., 2012). This complexity was further increased in RIM1α^- /-^. In view of the observed changes in AZ dimensions, molecular interactions of the multidomain protein RIM1α should be considered. Several RIM1α domains bind to VGCCs and its Zinc finger domain to SVs, therefore RIM is an essential link (Kaeser et al., 2011; Schoch et al., 2002). In addition, RIM1α interacts with core AZ components like ELKS/CAST, RIM-BP and liprin generating a multifunctional scaffold (Emperador-Melero and Kaeser, 2020). Therefore, the absence of RIM1α results in a disassembly of AZ scaffold leading to an increase in AZ area and variability. Although our EM tomography technic does not allow molecular mapping of individual proteins such as VGCCs and the above-named scaffold proteins in AZs, it appears likely that the observed increase in AZ area in accompanied by a redistribution of these components.

Since Castillo et al. (Castillo et al., 2002) mentioned no difference in AZ profile length in MFBs between RIM1α^-/-^ and WT mice, the larger AZ area and the increase in synaptic cleft height in the RIM1α^-/-^ was unexpected. While the absolute difference in cleft height between wildtype and RIM1α^-/-^ was only 2nm, the relative difference is 20%. However, the 20% change in height (2D) translates in much bigger cleft volume change in 3D which is likely to be of relevance for synaptic transmission. It is clear, that measurements in this range require excellent resolution. It would be interesting to study synaptic cleft height with our approach in other RIM knockout models such as of *Drosophila* (Graf et al., 2012). Finally, this finding should be viewed in context to the complex cleft ultrastructure (Martinez-Sanchez et al., 2021) and transsynaptic nanocolumns or indirect VGCC interactions with postsynaptic receptors (Tang et al., 2016) or with synaptic adhesion molecules, such as neurexin-1 (Brockhaus et al., 2018).

### Number and position of SVs are influenced by RIM1α

We report substantial reduction of (tight) SV docking at giant MFB AZs in the absence of RIM1α. This supports the role of RIM in SV docking/priming and is consistent with findings in synaptosomal preparations (Fernandez-Busnadiego et al., 2013) and in dissociated hippocampal cultures of RIM1/2 deficient mice (Zarebidaki et al., 2020). Tight SV docking is assumed to be mediated by an interaction of the Zinc finger of RIM with the C_2_A domain of Munc13-1 (Imig et al., 2014; Neher and Brose, 2018). For the Zinc finger of RIM, it is proposed, that it recruits Munc13 to the AZ (Andrews-Zwilling et al., 2006). Deletion of RIM1α lead to a 60% reduction of Munc13-1 expression (Andrews-Zwilling et al., 2006; Schoch et al., 2002). Mechanistically, the RIM Zinc finger is required for disruption of autoinhibitory homodimerization by forming a heteromeric complex with Munc13 (Deng et al., 2011). Thus, the reduced SV docking phenotype in RIM1α^-/-^ can be explained by these changes. Recent EM quantifications of SV docking associated the heterodimer of RIM/Munc13-1 with SV docking due to the formation of monomeric priming-competent Munc13 molecules (Camacho et al., 2017). A second explanation for reduced tight docking could be the reduction of the RIM C_2_B domain interaction with the phospholipid PIP_2_ in the AZ membrane, therefore disrupting the tight link between SVs and the release site (de Jong et al., 2018). Another key finding of our study is the delocalization of tightly docked SVs (0-2 nm) from the AZ c.o.m. in RIM1α^-/-^ (Figure 4). Regarding this observation, one may wonder about the molecular arrangement of release site defining AZ components such as Munc13-1 and VGCCs in RIM1α^-/-^. Interestingly, STED imaging in RIM-BP2 KO mice revealed, that the distance of RIM and Cav2.1 increases up to 35 % in MFBs (Brockmann et al., 2020). Electron tomographic analyses of NMJs in *C. elegans* have shown that in the absence of the RIM homologue *unc-10* SVs delocalize from dense projections, defining the center of the AZ, presumably due to a lost collaboration with *syd-2*, a homologue of the scaffold AZ protein α-Liprin (Stigloher et al., 2011).

### Electron dense material below the SV is altered in RIM1α^-/-^

We found EDM width reduced from 19.4 nm to 17.9 nm in tightly docked SVs (0-2 nm) in RIM1α^-/-^ AZs (Figure 3E). In addition, EDM width of SVs at 2.1-5 nm was further reduced to 14 nm in both genotypes. These findings may be related to recently described hexagonal Munc13-1 protein densities in cryo-EM (Grushin et al., 2022). The authors described a model with three states of Munc13-1 oligomers: In state 1, Munc13-1 is in the upright configuration under captured SVs of unassembled SNAREs. The closed hexagonal cage of pre-primed SVs in state 2 has a reduced diameter. In state 3 of primed SVs with approximate half-zippered SNARE pins the diameter of the hexagon widens because of calcium influx and binding to Munc-13 C_2_B domains. In general terms the magnitude and direction of the EDM changes in our study are in accordance with this 3-state model. Furthermore, the smaller EDM width in RIM1α^-/-^ might derive from the reduced expression of Munc13-1 at AZs (Grushin et al., 2022). An additional explanation might be the missing interaction of PIP_2_ with the C_2_B domain of RIM1α. Regarding SV size, we found a small but significant increase of the absolute SV diameter in RIM1α^-/-^ (Figure 3F). Interestingly, Imig et al. described a similar increase in SV diameter in Munc13-1/2 double knockout (Imig et al., 2014). However, the causal relation between the absence of Munc-13 or RIM1α and the enlargement of SVs remains unclear.

### Organization of the SV pool up to 200 nm is changed in RIM1α^-/-^

Consistent with previous findings (Fernandez-Busnadiego et al., 2013; Schoch et al., 2002), we report a reduced SV pool in proximity to the AZ membrane (0-50 nm, ∼ one SV diameter plus SV protein coverage zone) for RIM1α^-/-^. In addition, our study shows an overall reduction of the total SV pool within 200 nm (∼ 4 SV diameter) in the absence of RIM1α, suggesting two distinct ‘net’-works of neighboring SVs: a wider proximal pool (RRP) (Rizzoli and Betz, 2004; Rollenhagen and Lübke, 2010) and a denser cloud of SVs with increasing distance to the membrane without any differences in their nearest neighbor configuration (releasable pool) (Rizzoli and Betz, 2004; Rollenhagen and Lübke, 2010). Our differentiation is based on a 3D investigation of nearest neighbor distances. Further theories involving RIM, RIM-BP and VGCCs support the structural differentiation of the SV pool in functionally distinct parts (Milovanovic et al., 2018; Wu et al., 2019). Studies in conditional RIM/ELKS and ELKS1α/2α mice suggest that the interplay of both scaffold proteins is essential for presynaptic AZ composition including the SV pool size and that in particular ELKS control the size of the RRP through its N-terminal coiled-coil domains (Held et al., 2016; Wang et al., 2016). We found a more heterogeneous distribution of SVs within the SV pool up to 200 nm resulting in overall decreased SV density per area, however (and somehow counterintuitive) also decreased NN1-5 distances in RIM1α^-/-^ (Figure 5B, E, F). We believe this can be interpreted as accumulation of SVs in ‘nests’ in RIM1α^-/-^ compared to a more homogenous SV distribution in wildtype AZs. Although, the molecular mechanism of this ‘nest’ formation remains unclear, one might speculate that it relates to synapsins and their role in SV reserve pools (Zhang and Augustine, 2021).

We find an increased accumulation of mitochondria in RIM1α^-/-^ AZs (Figure S2). The expansion of AZ area, synaptic cleft and delocalization of docked SVs in RIM1α^-/-^ might raise energy consumption (Devine and Kittler, 2018; Pulido and Ryan, 2021). Greater proximity of mitochondria and AZs should facilitate calcium and ATP supply for the SV pool and the AZ membrane.

The complex geometry of the giant MFB AZs and its docked SV pool implicates that further studies are needed to fully decipher the molecular 3D nanoarchitecture of AZs as well as SV pool engrams as memory storage for short-term information (Vandael et al., 2020). More detailed information can be obtained by using advanced electron tomography (e.g. STEM tomography) for 3D mapping of entire AZs. This will help to interpret and correlate 3D AZ maps with functional measurements in acute brain slices.

## Acknowledgements

The authors thank G. Krohne (Würzburg), Y. Schwab and team (EMBL, Heidelberg) for fruitful discussions, further, D. Bunsen, C. Gehrig-Höhn and B. Trost for excellent technical assistance. This work has been supported by funding of the Deutsche Forschungsgemeinschaft (DFG, German Research Foundation) to M.H. and A.-L.S. (CRC 166, Project B06) and by the Interdisciplinary Clinical Research Center (IZKF) Würzburg to M.M.P. (Z-3/69), M.H. (N-229) and A.-L.S. (N-229). The JEOL JEM-2100Transmission Electron Microscope is funded by the Deutsche Forschungsgemeinschaft (DFG, German Research Foundation) – 218894163.

## Author contributions

*Conceptualization*: K.L., C.S., M.H. and A.-L.S.; *Methodology*: K.L., C.S., M.H., A.- L.S.; K.L. and M.P. established rapid cryo-immobilization of acute brain slices. S.S. provided the mouse line. *Investigation*: K.L., C.S.; *Formal analysis*: K.L. under supervision of M.M.P., P.K., C.S., M.H., A.-L.S.; *Validation, data interpretation*: K.L., M.M.P., C.S., M.H., A.-L.S.; *Software*: K.L., P.K.; *Resources*: C.S., M.H., A.-L.S.; *Funding acquisition*: M.M.P., C.S., M.H., A.-L.S.; *Visualization*: K.L., M.M.P.; *Data curation*: K.L., M.M.P., M.P., C.S., M.H., A.-L.S.; *Writing - original draft*: K.L., M.M.P., M.H., A-L.S.; *Writing - review & editing*: All authors. All authors read and approved the final manuscript.

## STAR Methods Animals

Three adult RIM1α knock-out mice (RIM1α^-/-^, C57BL/6 (Schoch et al., 2002) and three wild-type littermates (RIM1α^+/+^, C57BL/6) at an age of 13-19 weeks were used for the experiments. The procedures were carried out in accordance with the German regulations and guidelines for animal experimentation, the EU Directive 2010/63/EU as well as the United States Public Health Service’s Policy on Humane Care and Use of Laboratory Animals and approved by the district government of Lower Franconia in Germany as the responsible authority (Permit Number RUF-55.2.2.-2532-2-571-11).

### Slice preparation

Acute brain slices were prepared as described previously for patch-clamp recordings of hippocampal mossy fiber synapses (Bischofberger et al., 2006), with modifications. In brief, mice were anesthetized with isoflurane (1 mg/ml, CP-Pharma) and decapitated in deep anesthesia with a pair of scissors at the level of the cervical medulla. The head was dropped into ice-cold artificial cerebrospinal fluid (ACSF, containing 125 mM NaCl, 25 mM NaHCO_3_, 2.5 mM KCl, 1.25 mM NaH_2_PO_4_, 25 mM glucose, 2 mM CaCl_2_ and 1 mM MgCl_2_, equilibrated with 95% O2/5% CO2, pH 7.3, kept at -4°C). The skin surrounding the head was removed carefully, then the skull cap was opened with a single sagittal cut from the foramen magnum to the olfactory bulb. Right and left part of the skull cap were separated without any contact to the ventral side of the brain. Two coronal cuts were made at the level of cerebellum and olfactory bulb to separate the entire brain from its skull basis using a small spatula. Due to possible time sensitive changes of cellular brain morphology (Bischofberger et al., 2006), the procedure was kept at a maximum preparation time of 60-90 seconds. With its ventral surface, the brain was mounted on a cutting chamber, containing a high sucrose cutting solution (ACSF, supplemented with 75 mM sucrose), of a Leica VT1200 vibratome (Leica Microsystems). Four consecutive horizontal sections of 200 µm thickness were cut at a defined level of the dorsal hippocampus. This region was defined by the Allan Mouse Brain Atlas (Version: Mouse, Adult, 3D coronal) approximately 2600 µm below the dorsal brain surface) and beginning from the first clear appearance of both dentate gyrus (DG) and *cornu ammonis* (CA).

### Cryo-immobilization of acute brain slices

Cryo-immobilization of acute brain slices was conducted as described previously for morphological analysis of synaptic contacts in *C. elegans* and *M. musculus* (Borges-Merjane et al., 2020; Markert et al., 2020; Stigloher et al., 2011), with modifications. In brief, brain slices were transferred onto pre-cooled petri dishes coated with a silicone elastomer (Sylgard 184, Sigma-Aldrich) containing ice-cold ACSF. Slices were cut down to the left, entirely intact hippocampal region using a fine scalpel and loaded into high pressure freezing carriers (type A, 3 mm diameter and 200 µm depth, Leica Microsystems). The bottom carrier was overfilled slightly with precooled 15 % polyvinyl-pyrrolidone in ice-cold ACSF, so that the liquid formed a convex hull upon the carrier itself. To avoid harming tissue integrity during manipulation, slices were enclosed into ACSF droplets for transfer. A second carrier (type B, 3 mm diameter and 300 µm depth, Leica Microsystems) was placed onto the first serving as a lid, taking advantage of the convex hull formed by ACSF to avoid a formation of air bubbles within the final carrier sandwich. Samples were processed at a freezing speed > 20000 Ks^-1^ and a pressure > 2100 bar with the high-pressure freezing machine EM HPM100 (Leica Microsystems).

### Freeze substitution

High-pressure frozen brain samples were transferred separately into a single small Teflon container of the EM AFS2 freeze substitution (FS) system (Leica Microsystems). High pressure freezing chamber sandwiches were opened to allow better penetration during freeze substitution. Samples were incubated as described previously (Markert et al., 2020; Stigloher et al., 2011). Briefly, samples were kept in anhydrous acetone containing 0.1% tannic acid and 0.5% glutaraldehyde for 20h at - 90°C. Then, the solution was replaced entirely by a fresh solution to avoid possible hydrous saturation of acetone caused by the initial sample loading of the AFS system. Samples were kept in the first FS solution for a total of 96h at -90°C, followed by four washing steps with anhydrous acetone within 1h. Thereafter, the samples were fixated in 2% OsO_4_ in anhydrous acetone for 28h at -90°C and heated slowly by an increase of temperature from -90°C to -20°C within 14h. At -20°C, the pellets were incubated for 16h. Then, the temperature was increased to 4°C in 4h. After four washing steps (0.5h interval) with anhydrous acetone, the temperature was increased to 20°C in 1h. Subsequently, the samples were transferred in a freshly prepared epoxy resin solutions of 50% of the epoxy resin (EPON, component A: 50ml 2-dodecenylsuccinate anhydride, 31 ml EPON812; component B: 44.5 ml methyl-5-norbornene-2,3- dicarboxylic anhydride, 50 ml EPON812; component C: 0.5 ml 2,4,6-trisdimethylaminomethylphenole; SERVA Electrophoresis GmbH) in acetone for 3h at room temperature, 90% epoxy resin in acetone overnight at 4°C, and 3 times in 100% epoxy resin at room temperature. During the infiltration process, the intact tissue pellets were solved carefully from the carrier to allow better epoxy resin penetration.

### Flat embedding

Single tissue pellets, covered in 100% epoxy resin, were placed separately onto a sheet of transparent fluorinated-chlorinated thermoplastic (0.19812 mm thickness, ACLAR®, Ted Pella). Thereafter, a second thermoplastic sheet was positioned as a lid upon the sample, thereby avoiding air bubbles between both sheets. The sandwich-like ensemble was weighted down by a small handmade aglet and masked at its sides to avoid a leakage of epoxy resin. The samples were polymerized for 48h at 60°C.

### Sample processing

The flat-embedded hippocampal slices were glued onto epoxy resin nibs without a thermoplastic sheet in between using cyanoacrylate. Tissue blocks were trimmed manually to trapezia containing the whole CA3 region and characteristic parts of the granule cell (GC) region for further orientation. Both ultra-thin sections of 60-70 nm and semi-thin sections of 250 nm were cut with a Histo diamond knife (Diatome AG) at an EM UC7 ultramicrotome (Leica Microsystems). Ultra-thin sections were positioned onto Pioloform (5% polyvinyl butyral in trichloromethane) coated copper grids (50 mesh, G2050C, Plano GmbH) to evaluate high pressure freezing quality of the tissue. 4-5 semi-thin hippocampal sections were positioned serially onto Pioloform coated, single slotted copper grids (2 × 1 mm, G2500C, Plano GmbH) for electron tomography. To prevent electron charging particularly at high tilt angles, single slot grids were coated by an approximately 3 nm thin layer of carbon using the high-vacuum carbon coating unit CCU-010 (Safematic GmbH). Both types of sections were contrasted with 5% uranyl acetate in ethanol for 7.5 min and 50% Reynolds’ lead citrate (Reynolds, 1963) in ddH_2_O for 10 min. In between both steps, the samples were washed in pure ethanol, in 50% ethanol in ddH_2_O and pure ddH_2_O; after the contrasting, samples were washed three times in ddH_2_O and blotted dry with filter paper.

### Image acquisition

Both, electron micrographs and tilt image series of synapses were acquired at 200 kV using a JEM-2100 (JEOL) electron microscope equipped with a TemCam F416 4k×4k camera (Tietz Video and Imaging Processing Systems). Tilt series acquisition was carried out within a minimum range of −60° to +60° tilt angle with 1° increments at a pixel size of 0.287 nm. The SerialEM software package (Mastronarde, 2005) was used for image acquisition. To ensure an acquisition of presynaptic active zones (AZ) with an extraordinary AZ profile length, especially in the KO phenotype, the resolution had to be adjusted to 0.3897 nm per pixel at selected synapses during the experiments. For optimal acquisition conditions, a full electron beam alignment was conducted prior to each tilt series.

### Selection of samples and region of interest

All HPF/FS, epoxy resin embedded brain slices were evaluated depending on possible freezing and/or embedding artefacts, i.e., formation of ice crystals (in particular, in the chromatin of cell nuclei, where they typically appear first), volumetric changes of cellular compartments and extracellular space, the appearance and intactness of lipid double layers, and the preservation of characteristic intracellular electron dense structures such as mitochondria and actin filaments. All samples were evaluated in random order, coded by sample embedding number and not phenotype. Only tissue slices which fulfilled the quality criteria for HPF/FS tissue (Weimer, 2006) were further processed for electron tomography. We observed that the quality of the HPF/FS brain sections declined depending on the time point of cutting at the vibratome. This may underpin a time sensitivity of the described protocol. Further, electron tomographic semi-thin sections were obtained within a minimum distance of 20-40 µm to the surface in slices with optimal tissue quality. If so, we could not observe any increased alteration of morphology due to possible cutting artefacts within 100 µm to the surface of the brain slice (Rostaing et al., 2006), or increasing formation of ice crystals with increasing distance to the center of the brain slice (Siksou et al., 2007). We collected tilt series at synapses on intact 2-5 µm giant mossy fiber boutons alongside the left supra-pyramidal mossy fiber tract (*Stratum lucidum*) of the CA3b region (Masukawa et al., 1982). Spine heads of CA3b pyramidal neurons had to be clearly identifiable by both spine apparatus (Gray, 1959), and postsynaptic filaments (PSF). AZ membranes were defined by corresponding PSFs connected to the postsynaptic membranes at the spine heads. Membrane parts with and without PSFs were clearly distinguishable. Beside synaptic contacts at MF-CA3 pyramidal spine heads, all other synaptic contacts within the MFB were excluded from further analyses. To outweigh individual heterogeneity of MFB presynapses (Pauli et al., 2021; Wilke et al., 2013), we aimed to analyze a minimum of 10 reconstructed tomograms per animal. Only tomograms in which both pre- and postsynaptic membranes were clearly identifiable throughout the entire z stack were included into analysis.

### Image analysis and tomogram segmentation

Tilt-image series alignment and tomographic reconstruction was performed with the ETomo/IMOD software package using a batch processing alignment and weighted back projection algorithm (Kremer et al., 1996). Overall, we observed a better alignment of images and final tomogram quality with the applied reconstruction protocol of systematically placed patches in high resolution compared to a reconstruction based on gold fiducials. Further, a fiducial free preparation was preferred to minimize possible artefacts due to gold fiducials within the AZ scene in tilt series of high magnification. All reconstructed tomograms were randomized by random number sampling prior to segmentation and 3D reconstruction. Segmentation and 3D reconstruction of electron tomograms were carried out in full resolution without binning using the 3Dmod software of the IMOD software package (Kremer et al., 1996). All types of synaptic vesicles (SVs) were annotated as ideal spherical objects by setting a point at the vesicle center in virtual section with its largest vesicle diameter using the drawing tool ‘normal’. Further, the point was resized to the extent of the outer vesicle membrane using the mouse wheel in the drawing tool ‘normal’. Within the reconstructed tomograms, we observed that SVs often deviate from an ideal spherical object and, despite possible deformation, were clearly distinguishable from smooth endoplasmatic reticulum. SVs of 0-200 nm distance to the presynaptic membrane were included into the annotation and defined as SV pool (Rollenhagen et al., 2007). SVs which were tethered or docked within the peri-synaptic area (lateral distance of 200 nm to AZ edge on the same virtual section, cf. mean AZ spacing at giant MFB AZs in rats: 0.40 (P28) - 0.48 µm (adult) (Rollenhagen et al., 2007) and with a distance up to 10 nm to the plasma membrane were annotated as vesicles connected outside of the AZ. Presynaptic membranes were annotated as ‘open’ line objects, of which gaps of 15-20 virtual sections were linearly interpolated using the tool ‘interpolator’. The synaptic cleft volume was quantified via one or more separate ‘closed’ objects.

Every 20 virtual section, the area between the inner part of the double lipid membranes of pre- and postsynapse was annotated and interpolated in between using the tool ‘interpolator’ and the ‘smooth’ function. If needed, contours were manually corrected after interpolation. The extent of the synaptic cleft was defined by the extent of postsynaptic filaments. To improve the illustration of SVs in the 3D tomograms, the following settings were uniformly used: grade 4 in global quality of points, option ‘fill’ in the ‘drawing style’.

### 2D and 3D quantitative data analyses

3D coordinates of SV centers and membranes, vesicle radii, and lengths of open and closed ‘line’ objects were extracted via command line using ‘imodinfo’ and ‘model2point’. Area and volume information were assessed by using the ‘mesh’ function. To avoid inaccurate mesh calculation due to complex branched AZ shapes, sub-meshes of presynaptic membranes and the synaptic cleft were calculated separately as distinct surfaces. The distance of SVs (0-10 nm from AZ membrane) and electron dense material connecting SV and presynaptic membrane was manually measured in 3Dmod. The main width of electron dense material (EDM) connecting SV and membrane was measured at the virtual slice of the defined SV center. The existence of a main EDM was observed in nearly all SVs between 0-10 nm in the entire data set. SVs with a distance of 0-2 nm to the presynaptic membrane were defined as docked. To avoid measuring artefacts, the ‘slicer’ window function was used to visualize the entire contact zone between SV and presynaptic membrane. 3D coordinates were processed via text files in Jupyter Notebook 6.2.0 (Anaconda), a web-based programming environment for Python (van Rossum and de Boer, 1991), Version 3.7, with a custom-written script. Stereogeometrical data were calculated using ‘numpy’ (Harris et al., 2020), ‘scikit-learn’ (Pedregosa et al., 2011) and ‘shapely’ (Gillies et al., 2007). Mathematical calculations are described in detail in supplemental Python scripts. For 2D quantification of mitochondria within 0-500 nm distance to giant MFB AZ membranes and their perisynaptic membranes, a randomly placed systematic grid lattice with a point spacing of 22.36 nm was used as an overlay on electron micrographs of equal magnification (15,000x). Points were counted depending on their localization (‘M’ = mitochondria, ‘NM’ = no mitochondria) and categorized in 5 bands of 100 nm distance each to the membranes (0-100 nm, 100-200nm etc.). 2D AZ profiles of RIM1α^-/-^ giant MFBs (N = 45) and controls (N = 20) were selected upon a clear presence of mitochondria near the AZ profile and excellent morphological ultrastructure of the giant MFB. In case of several AZs at a spine head, the AZ nearest to the mitochondrial accumulation was quantified. In case of the quantification at the perisynaptic membrane, mitochondria were only quantified in case they were clearly associated to the depicted one AZ profile and not ambiguously to nearby spaced AZ profiles at the same spine head. The present method for 2D spatial stereology was modified from C. Hacker and J. Lucocq (Hacker and Lucocq, 2014).

### Statistics and visualization

Statistical analyses were carried out with the statistical software programs Sigma Plot 14 (Systat Software GmbH) and OriginPro 2021 (Origin Lab). 2D stereology of mitochondria was carried out in Fiji Version v1.53n (Schindelin et al., 2012). The number of animals and the sample size for tomograms was defined *a priori* according to standard publications in electron tomography (Imig et al. 2014). The experimental procedure was independently replicated several times within the laboratory. For all data, normality was tested using Shapiro-Wilk tests (sample size N ≤ 5000) and Kolmogorov-Smirnov-tests. Non-parametric data were analyzed using non-parametric Mann-Whitney rank sum tests. Parametric data were analyzed using Welch’s t-tests. The power (PW) of conducted analyses is reported separately. For both groups, non-equal variances were assumed due to the highly experimental character of observed (rare) phenomena (e.g., ‘tightly’ docked SVs). Correlations were calculated using the Spearman rank order correlation. Comparison of data distributions were calculated using the Akaike information criterion and the F-test. P-values were calculated with a minimum of 6 decimal places, p-values with a probability < 0.0001 are reported as rounded values. Data are reported as median ± 25^th^ and 75^th^ percentile for non-parametric data, unless indicated otherwise, and as mean ± SD for parametric data. Illustrator 2021 (Adobe Creative Cloud) was used for further illustration.

## Data availability

The datasets and computer codes produced in this study will be available upon request and after publication online.

## Supplemental Information

**Supplemental Figure S1.**
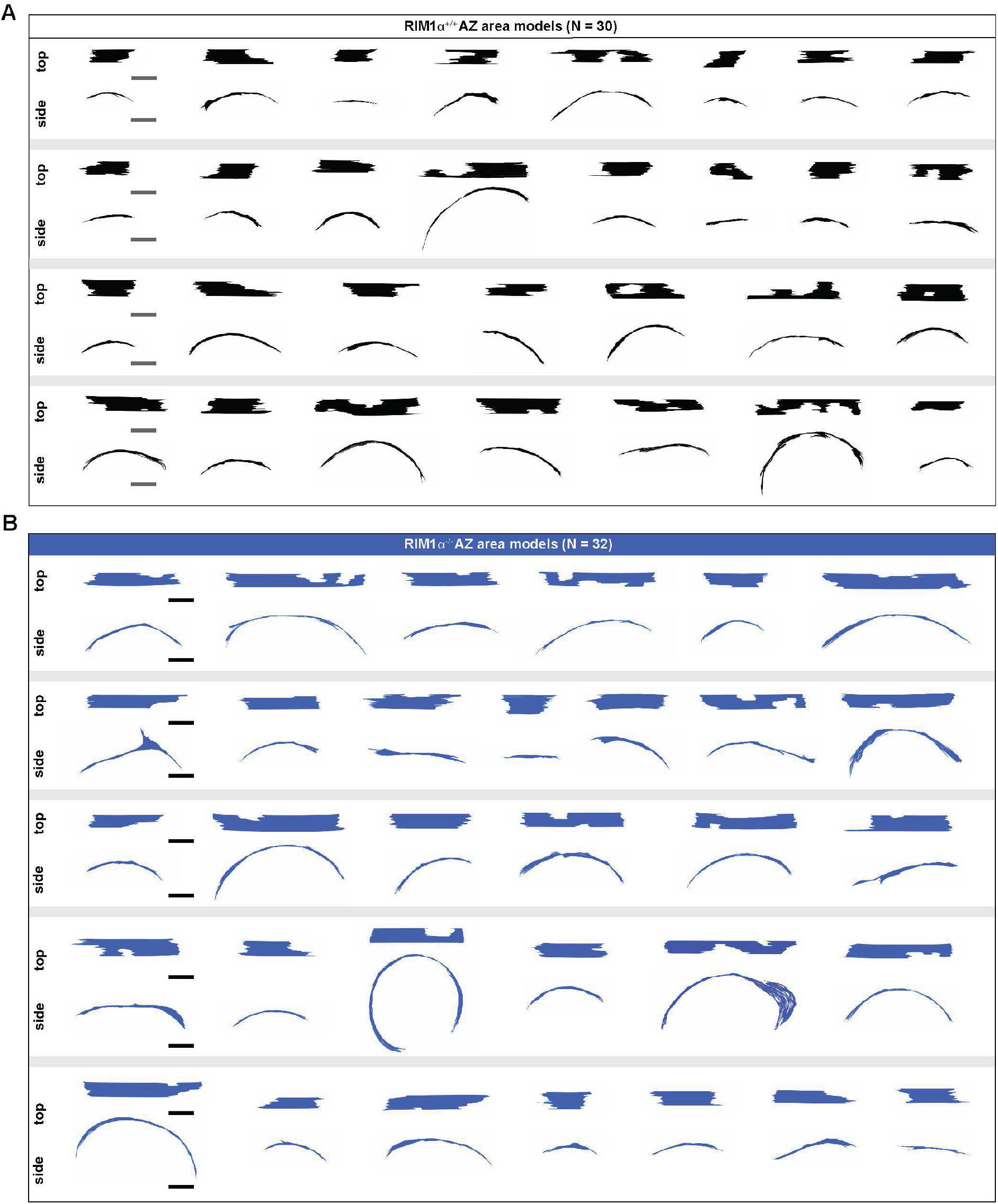
IMOD models of presynaptic giant MFB AZ membranes in RIM1α^+/+^ and RIM1α^-/-^ mice. **A** Presynaptic AZ membranes of electron tomograms illustrated as annotated centered IMOD objects of RIM1α^+/+^ (black) mice. Illustration in top and side view. **B** IMOD objects of presynaptic RIM1α^-/-^ (blue) AZ membranes. Scale bars: (A, B) 100 nm.

**Supplemental Figure S2.**
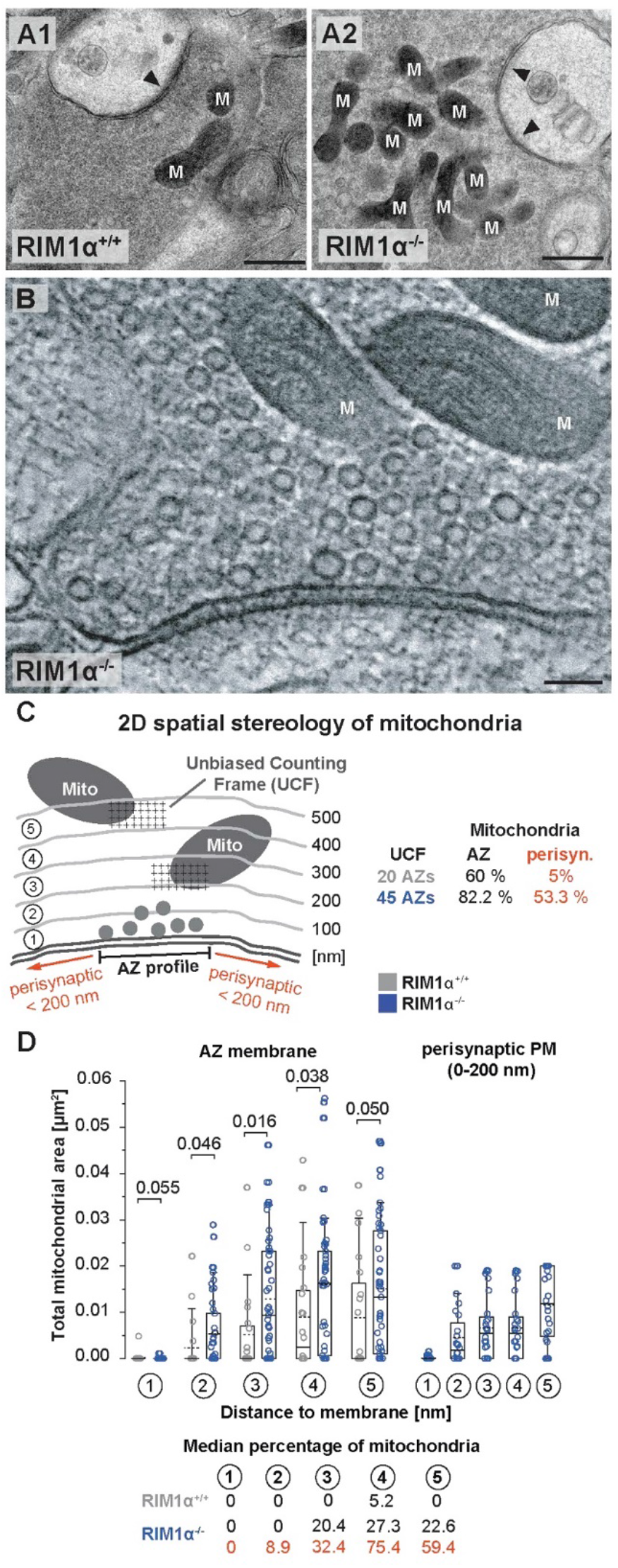
Mitochondria at giant MFB AZs. **A** Exemplary standardized electron micrographs of giant MFB AZs (arrowhead) and associated mitochondria (M) in RIM1α^+/+^ (A1) and RIM1α^-/-^ (A2) used for 2D stereology to estimate mitochondrial area near depicted AZs. **B** Exemplary electron tomographic slice of included RIM1α^-/-^ AZ tomogram illustrates the spatial proximity of mitochondria to the SV pool and the AZ. **C** An unbiased counting frame was used in 45 RIM1α^-/-^ AZ profiles and 20 controls to quantify mitochondrial area in bands of 100 nm distance to the membrane, both in relation to the AZ and the perisynaptic area up to 200 nm distance to both AZ edges (modified from Hacker et al. 2014). **D** RIM1α^-/-^ AZs show increased mitochondrial area within 500 nm distance to the AZ membrane. In the experiment, only RIM1α^-/-^ AZs had clearly associated mitochondria within 200 nm to the AZ edges compared to controls (1 of 20 AZ profiles).

**Supplemental Figure S3.**
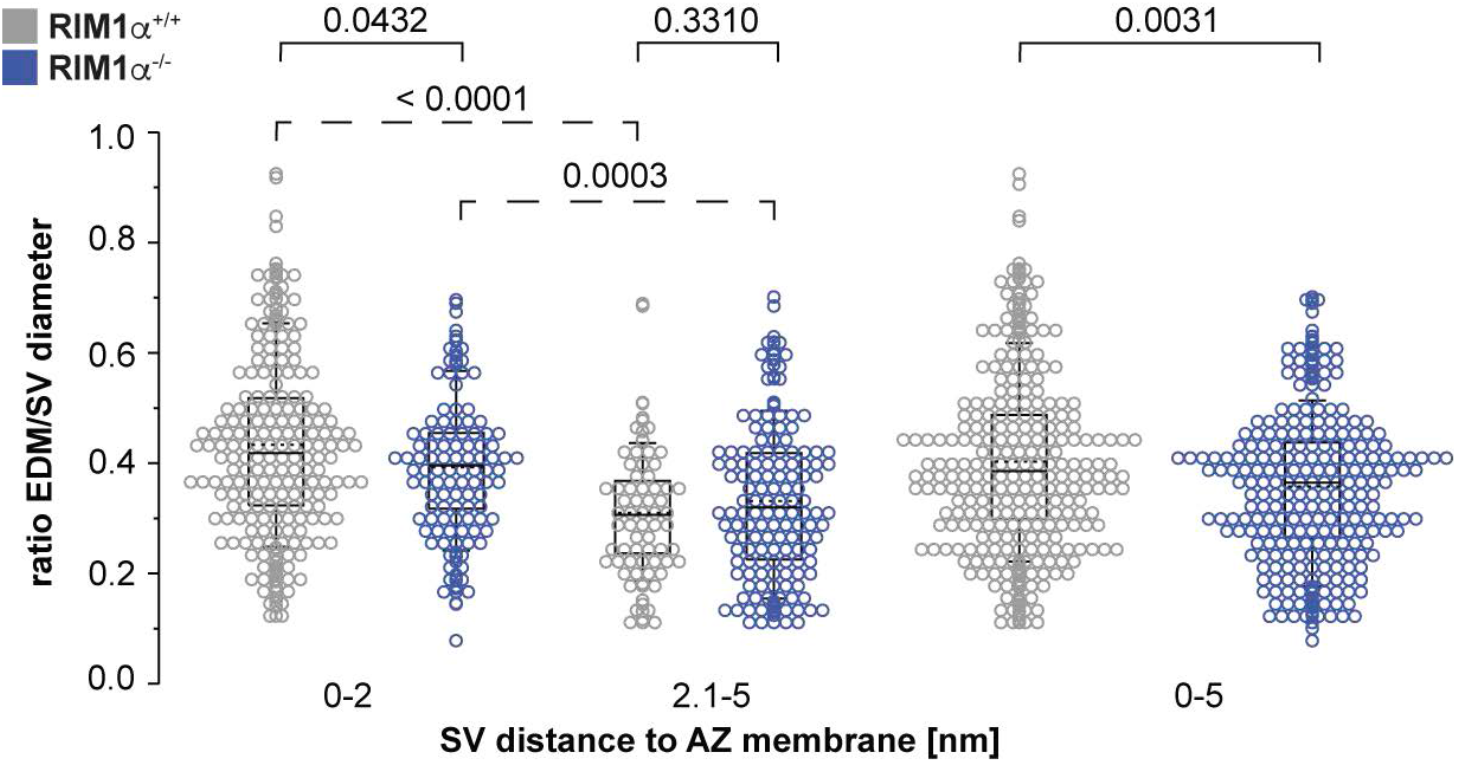
Ratio of EDM width and SV diameter in the docked SV pool at giant MFB AZs. Summary graphs of the maximal EDM width normalized to the SV diameter in 0-2 nm, 2.1-5 nm and 0-5 nm distance to the AZ membrane in both groups.

**Supplemental Videos SA1-SA4. Electron tomograms of docked synaptic vesicles in wild-type and RIM1α KO active zones in high magnification**.

**A1-2** Video file (.avi) of SVs near or at the presynaptic membrane in wild-type giant MFB to CA3b spine head synapses of two animals. Electron tomographic slices were extracted from original tomograms and processed as video file.

**A3-4** SVs near or at the presynaptic membrane in RIM1α deficient giant MFB to CA3b spine head synapses of two animals.

**Supplemental Table S1.** Summary of numerical and statistical values.

| Parameter | RIM1α<sup>-/-</sup> | RIM1α<sup>+/+</sup> | p-value | Statistics | Figure |
| --- | --- | --- | --- | --- | --- |
| **AZ area [$\mu\text{m}^2$]** | 0.100, 0.072-0.132 (25-75%), N = 32 AZs | 0.068, 0.048-0.098 (25-75%), N = 30 AZs | 0.0019 | MWRS | **2J** |
| **AZ profile length [nm]** | 513.49, 330.33-665.85, N = 12,750 | 338.77, 241.16-460.77, N = 13,991 | < 0.0001 | MWRS | **2K** |
| **max. AZ area extent [nm]** | 802.50, SD = 208.92, N = 32 | 602.31, SD = 159.14, N = 30 | < 0.0001, PW = 0.99 | Welch's test | **2L** |
| **synaptic cleft height [nm]** | 10.08, SD = 2.03, N = 32 AZs | 8.45, SD = 1.99, N = 30 AZs | 0.0023, PW = 0.88 | Welch's test | **2M** |
| **SVs/0.01 $\mu\text{m}^2$ AZ area** |  |  |  |  |  |
| **0-2 nm** | 0.341, 0.108-0.469, N = 32 AZs | 1.133, 0.674-1.410, N = 30 AZs | < 0.0001 | MWRS | **3B** |
| **0-5 nm** | 0.946, SD = 0.477, N = 32 AZs | 1.662, SD = 1.622, N = 30 AZs | < 0.0001, PW = 0.99 | Welch's test | **3B** |
| **2.1-5 nm** | 0.612, SD = 0.365, N = 32 AZs | 0.488, SD = 0.365, N = 30 AZs | 0.1231, PW = 0.34 | Welch's test | **3B** |
| **5-10 nm** | 0.157, 0.083-0.455, N = 32 AZs | 0.222, 0.071-0.455, N = 30 AZs | 0.8707 | MWRS | **3B** |
| **SV distance from AZ membrane [nm]** | 3.36, 1.96-5.10, N = 429 SVs | 1.75, 0.00-3.88, N = 428 SVs | < 0.0001 | MWRS | **3B** |
| **SV distance to PM [nm]** | 6.01, 3.75-8.12, N = 91 SVs | 3.47, 1.46-6.75, N = 86 SVs | < 0.0001 | MWRS | **3C** |
| **EDM 0-2 nm [nm]** | 17.87, SD 5.62, N = 101 SVs | 19.43 nm, SD 6.42, N = 188 SVs | 0.0331, PW = 0.57 | Welch's test | **3E** |
| **EDM 2.1-5 nm [nm]** | 13.86, 9.69-18.53, N = 136 SVs | 13.59 nm, 10.68-17.00, N = 62 SVs | 0.6649 | MWRS | **3E** |
| **SV diameter [nm]** | 43.96, 41.33-46.76, N = 4,079 SVs | 43.39, 41.33-46.15, N = 3,111 SVs | < 0.0001 | MWRS | **3F** |
| **distance of** |  |  |  |  |  |
| **NN1 [nm]** | 50.57, 31.00-101.83, N = 110 SVs | 39.05, 28.12-57.88, N = 225 SVs | < 0.0001 | MWRS | **4B** |
| **NN2 [nm]** | 111.32, 75.50-227.31, N = 110 SVs | 60.97, 42.90-89.15, N = 225 SVs | < 0.0001 | MWRS | **4B** |
| **NN3 [nm]** | 255.59, 136.25-374.15, | 89.42, 62.32-134.23, N = 216 SVs | < 0.0001 | MWRS | **4B** |
|  | N = 101 SVs |  |  |  |  |
| **NN4 [nm]** | 310.94, 194.49-443.03,<br>N = 89 SVs | 105.27, 81.53-159.60,<br>N = 203 SVs | < 0.0001 | MWRS | **4B** |
| **NN5 [nm]** | 310.94, 207.22-442.37,<br>N = 59 SVs | 141.10, 99.92-215.09,<br>N = 188 SVs | < 0.0001 | MWRS | **4B** |
| **distance SV center to AZ edge 1 [nm]** | 404.65, 186.92-632.44,<br>N = 108 SVs | 219.07, 120.55-341.93,<br>N = 203 SVs | < 0.0001 | MWRS | **4D** |
| **distance SV center to AZ edge 2 [nm]** | 550.76, 299.09-915.65,<br>N = 108 SVs | 227.59, 107.09-381.22,<br>N = 203 SVs | < 0.0001 | MWRS | **4D** |
| **AZ edge 1 to AZ edge 2 distance ratio** | 0.389, 0.207-0.643 to 0.611, 0.357-0.793 | 0.479, 0.251-0.744 to 0.521, 0.256-0.748 | 0.0310 | MWRS | **4E** |
| **Relative distance AZ edge 1 vs. AZ edge 2** | 0.389, 0.207-0.643 vs. 0.611, 0.357-0.793 |  | < 0.0001 | MWRS |  |
| **Relative distance AZ edge 1 vs. AZ edge 2** |  | 0.479, 0.251-0.744 vs. 0.521, 0.256-0.748 | 0.8590 | MWRS |  |
| **distance of docked SV to 3D c.o.m. [nm]** | 195.03, 118.28-312.79,<br>N = 108 | 110.59, 68.73-187.13,<br>N = 203 | < 0.0001 | MWRS | **4F** |
| **correlation of SV pool size (0-200 nm) with AZ area** | r = 0.721,<br>N = 32 AZs | r = 0.829,<br>N = 30 AZs | RIM1$\alpha^{-/-}$:<br>< 0.0001<br>RIM1$\alpha^{+/+}$:<br>< 0.0001 | SROC | **5A** |
| **correlation of SV pool size (0-50 nm) with AZ area** | r = 0.611,<br>N = 32 AZs | r = 0.871,<br>N = 30 AZs | RIM1$\alpha^{-/-}$:<br>< 0.0002<br>RIM1$\alpha^{+/+}$:<br>< 0.0001 | SROC | **5A** |
| **comparison of SV pool (0-50 nm)/AZ area distribution** | see before | see before | 0.0066 (F = 5.4892), | F-test |  |
| **total SV pool [SVs/0.01 $\mu\text{m}^2$]** | 12.14, SD = 3.33, N = 32 AZs | 15.26, SD = 4.24, N = 30 AZs | 0.0022, PW = 0.86 | Welch's test |  |
| ***SVs/0.01 $\mu\text{m}^2$ AZ area*** |  |  |  |  |  |
| **0-50 nm** | 2.70, SD 0.79 | 3.66, SD 0.89 | < 0.0001, PW = 0.99 | Welch's test | **5B** |
| **51-100 nm** | 3.23, SD 1.01 | 3.82, SD 1.20 | 0.0434, PW = 0.53 | Welch's test | **5B** |
| **101-150 nm** | 3.54, SD 1.17 | 4.24, SD 1.40 | 0.0371, PW = 0.56 | Welch's test | **5B** |
| **151-200 nm** | 2.69, SD 0.95 | 3.51, SD 1.32 | 0.0069, PW = 0.79 | Welch's test | **5B** |
| ***NN distance [nm] (SV pool 0-50 nm)*** |  |  |  |  |  |
| **NN1** | N = 964 SVs<br>27.90, 23.63-33.02 | N = 741 SVs<br>29.52, 24.55-34.35 | 0.0035 | MWRS | **5E** |
| **NN2** | 34.15, 29.63-40.95 | 35.50, 31.42-41.46 | 0.0030 | MWRS | **5E** |
| **NN3** | 39.69, 33.99-48.15 | 40.86, 35.61- 48.31 | 0.0376 | MWRS | **5E** |
| **NN4** | 45.62, 38.45-56.11 | 45.80, 39.20-55.34 | 0.8196 | MWRS | **5E** |
| **NN5** | 51.892, 43.21-63.43 | 51.86, 43.75-61.57 | 0.7291 | MWRS | **5E** |
| ***NN distance [nm] (SV pool 51-200 nm)*** |  |  |  |  |  |
| **NN1** | N = 3,256 SVs<br>26.00, 22.36- 30.04 | N = 2,443 SVs<br>27.88, 24.37-31.60 | < 0.0001 | MWRS | **5F** |
| **NN2** | 30.95, 27.70-35.10 | 32.77, 29.12-36.78 | < 0.0001 | MWRS | **5F** |
| **NN3** | 34.83, 30.96-39.64 | 36.42, 32.53-40.86 | < 0.0001 | MWRS | **5F** |
| **NN4** | 38.39, 33.89-44.52 | 39.65, 35.46-45.47 | < 0.0001 | MWRS | **5F** |
| **NN5** | 42.16, 36.67-49.83 | 43.18, 38.19-50.44 | < 0.0001 | MWRS | **5F** |

## References

Andrews-Zwilling, Y.S., Kawabe, H., Reim, K., Varoqueaux, F., and Brose, N. (2006). Binding to Rab3A-interacting molecule RIM regulates the presynaptic recruitment of Munc13-1 and ubMunc13-2. The Journal of biological chemistry 281, 19720–19731.

Bischofberger, J., Engel, D., Li, L., Geiger, J.R., and Jonas, P. (2006). Patch-clamp recording from mossy fiber terminals in hippocampal slices. Nat Protoc 1, 2075–2081.

Borges-Merjane, C., Kim, O., and Jonas, P. (2020). Functional Electron Microscopy, “Flash and Freeze,” of Identified Cortical Synapses in Acute Brain Slices. Neuron 105, 992–1006 e1006.

Brockhaus, J., Schreitmuller, M., Repetto, D., Klatt, O., Reissner, C., Elmslie, K., Heine, M., and Missler, M. (2018). alpha-Neurexins Together with alpha2delta-1 Auxiliary Subunits Regulate Ca(2+) Influx through Cav2.1 Channels. The Journal of neuroscience : the official journal of the Society for Neuroscience 38, 8277–8294.

Brockmann, M.M., Zarebidaki, F., Camacho, M., Grauel, M.K., Trimbuch, T., Sudhof, T.C., and Rosenmund, C. (2020). A Trio of Active Zone Proteins Comprised of RIM-BPs, RIMs, and Munc13s Governs Neurotransmitter Release. Cell Rep 32, 107960.

Butola, T., Alvanos, T., Hintze, A., Koppensteiner, P., Kleindienst, D., Shigemoto, R., Wichmann, C., and Moser, T. (2021). RIM-Binding Protein 2 Organizes Ca(2+) Channel Topography and Regulates Release Probability and Vesicle Replenishment at a Fast Central Synapse. The Journal of neuroscience : the official journal of the Society for Neuroscience 41, 7742–7767.

Camacho, M., Basu, J., Trimbuch, T., Chang, S., Pulido-Lozano, C., Chang, S.S., Duluvova, I., Abo-Rady, M., Rizo, J., and Rosenmund, C. (2017). Heterodimerization of Munc13 C2A domain with RIM regulates synaptic vesicle docking and priming. Nat Commun 8, 15293.

Castillo, P.E., Schoch, S., Schmitz, F., Sudhof, T.C., and Malenka, R.C. (2002). RIM1alpha is required for presynaptic long-term potentiation. Nature 415, 327–330.

de Jong, A.P.H., Roggero, C.M., Ho, M.R., Wong, M.Y., Brautigam, C.A., Rizo, J., and Kaeser, P.S. (2018). RIM C2B Domains Target Presynaptic Active Zone Functions to PIP2-Containing Membranes. Neuron 98, 335–349 e337.

Deng, L., Kaeser, P.S., Xu, W., and Sudhof, T.C. (2011). RIM proteins activate vesicle priming by reversing autoinhibitory homodimerization of Munc13. Neuron 69, 317–331.

Devine, M.J., and Kittler, J.T. (2018). Mitochondria at the neuronal presynapse in health and disease. Nat Rev Neurosci 19, 63–80.

Emperador-Melero, J., and Kaeser, P.S. (2020). Assembly of the presynaptic active zone. Current opinion in neurobiology 63, 95–103.

Fernandez-Busnadiego, R., Asano, S., Oprisoreanu, A.M., Sakata, E., Doengi, M., Kochovski, Z., Zurner, M., Stein, V., Schoch, S., Baumeister, W., et al. (2013). Cryo-electron tomography reveals a critical role of RIM1alpha in synaptic vesicle tethering. J Cell Biol 201, 725–740.

Gillies, S., Bierbaum, A., Lautaportti, K., and Tonnhofer, O. (2007). Shapely: Manipulation and Analysis of Geometric Objects. Available online at: https://github.com/Toblerity/Shapely.

Goodsell, D.S., Olson, A.J., and Forli, S. (2020). Art and Science of the Cellular Mesoscale. Trends Biochem Sci 45, 472–483.

Graf, E.R., Valakh, V., Wright, C.M., Wu, C., Liu, Z., Zhang, Y.Q., and DiAntonio, A. (2012). RIM promotes calcium channel accumulation at active zones of the Drosophila neuromuscular junction. The Journal of neuroscience : the official journal of the Society for Neuroscience 32, 16586–16596.

Gray, E.G. (1959). Electron microscopy of synaptic contacts on dendrite spines of the cerebral cortex. Nature 183, 1592–1593.

Grushin, K., Kalyana Sundaram, R.V., Sindelar, C.V., and Rothman, J.E. (2022). Munc13 structural transitions and oligomers that may choreograph successive stages in vesicle priming for neurotransmitter release. Proc Natl Acad Sci U S A 119.

Hacker, C., and Lucocq, J.M. (2014). Analysis of specificity in immunoelectron microscopy. Methods Mol Biol 1117, 315–323.

Han, Y., Kaeser, P.S., Sudhof, T.C., and Schneggenburger, R. (2011). RIM determines Ca(2)+ channel density and vesicle docking at the presynaptic active zone. Neuron 69, 304–316.

Harlow, M.L., Ress, D., Stoschek, A., Marshall, R.M., and McMahan, U.J. (2001). The architecture of active zone material at the frog’s neuromuscular junction. Nature 409, 479–484.

Harris, C.R., Millman, K.J., van der Walt, S.J., Gommers, R., Virtanen, P., Cournapeau, D., Wieser, E., Taylor, J., Berg, S., Smith, N.J., et al. (2020). Array programming with NumPy. Nature 585, 357–362.

Held, R.G., Liu, C., and Kaeser, P.S. (2016). ELKS controls the pool of readily releasable vesicles at excitatory synapses through its N-terminal coiled-coil domains. eLife 5, e14862.

Henze, D.A., Wittner, L., and Buzsaki, G. (2002). Single granule cells reliably discharge targets in the hippocampal CA3 network in vivo. Nat Neurosci 5, 790–795.

Hibino, H., Pironkova, R., Onwumere, O., Vologodskaia, M., Hudspeth, A.J., and Lesage, F. (2002). RIM binding proteins (RBPs) couple Rab3-interacting molecules (RIMs) to voltage-gated Ca(2+) channels. Neuron 34, 411–423.

Imig, C., Min, S.W., Krinner, S., Arancillo, M., Rosenmund, C., Sudhof, T.C., Rhee, J., Brose, N., and Cooper, B.H. (2014). The morphological and molecular nature of synaptic vesicle priming at presynaptic active zones. Neuron 84, 416–431.

Jonas, P., Major, G., and Sakmann, B. (1993). Quantal components of unitary EPSCs at the mossy fibre synapse on CA3 pyramidal cells of rat hippocampus. J Physiol 472, 615–663.

Kaeser, P.S., Deng, L., Fan, M., and Sudhof, T.C. (2012). RIM genes differentially contribute to organizing presynaptic release sites. Proc Natl Acad Sci U S A 109, 11830–11835.

Kaeser, P.S., Deng, L., Wang, Y., Dulubova, I., Liu, X., Rizo, J., and Sudhof, T.C. (2011). RIM proteins tether Ca2+ channels to presynaptic active zones via a direct PDZ-domain interaction. Cell 144, 282–295.

Kaeser, P.S., and Regehr, W.G. (2017). The readily releasable pool of synaptic vesicles. Current opinion in neurobiology 43, 63–70.

Kheirbek, M.A., Drew, L.J., Burghardt, N.S., Costantini, D.O., Tannenholz, L., Ahmari, S.E., Zeng, H., Fenton, A.A., and Hen, R. (2013). Differential control of learning and anxiety along the dorsoventral axis of the dentate gyrus. Neuron 77, 955–968.

Kintscher, M., Wozny, C., Johenning, F.W., Schmitz, D., and Breustedt, J. (2013). Role of RIM1alpha in short- and long-term synaptic plasticity at cerebellar parallel fibres. Nat Commun 4, 2392.

Kiyonaka, S., Wakamori, M., Miki, T., Uriu, Y., Nonaka, M., Bito, H., Beedle, A.M., Mori, E., Hara, Y., De Waard, M., et al. (2007). RIM1 confers sustained activity and neurotransmitter vesicle anchoring to presynaptic Ca2+ channels. Nat Neurosci 10, 691–701.

Korogod, N., Petersen, C.C., and Knott, G.W. (2015). Ultrastructural analysis of adult mouse neocortex comparing aldehyde perfusion with cryo fixation. eLife 4, e05793.

Kremer, J.R., Mastronarde, D.N., and McIntosh, J.R. (1996). Computer visualization of three-dimensional image data using IMOD. J Struct Biol 116, 71–76.

Kusick, G.F., Chin, M., Raychaudhuri, S., Lippmann, K., Adula, K.P., Hujber, E.J., Vu, T., Davis, M.W., Jorgensen, E.M., and Watanabe, S. (2020). Synaptic vesicles transiently dock to refill release sites. Nat Neurosci 23, 1329–1338.

Markert, S.M., Skoruppa, M., Yu, B., Mulcahy, B., Zhen, M., Gao, S., Sendtner, M., and Stigloher, C. (2020). Overexpression of an ALS-associated FUS mutation in C. elegans disrupts NMJ morphology and leads to defective neuromuscular transmission. Biol Open 9, bio055129.

Martinez-Sanchez, A., Laugks, U., Kochovski, Z., Papantoniou, C., Zinzula, L., Baumeister, W., and Lucic, V. (2021). Trans-synaptic assemblies link synaptic vesicles and neuroreceptors. Sci Adv 7, eabe6204.

Mastronarde, D.N. (2005). Automated electron microscope tomography using robust prediction of specimen movements. Journal of Structural Biology 152, 36–51.

Masukawa, L.M., Benardo, L.S., and Prince, D.A. (1982). Variations in electrophysiological properties of hippocampal neurons in different subfields. Brain Res 242, 341–344.

Maus, L., Lee, C., Altas, B., Sertel, S.M., Weyand, K., Rizzoli, S.O., Rhee, J., Brose, N., Imig, C., and Cooper, B.H. (2020). Ultrastructural Correlates of Presynaptic Functional Heterogeneity in Hippocampal Synapses. Cell Rep 30, 3632–3643 e3638.

Miki, T., Kiyonaka, S., Uriu, Y., De Waard, M., Wakamori, M., Beedle, A.M., Campbell, K.P., and Mori, Y. (2007). Mutation associated with an autosomal dominant cone-rod dystrophy CORD7 modifies RIM1-mediated modulation of voltage-dependent Ca2+ channels. Channels (Austin) 1, 144–147.

Milovanovic, D., Wu, Y., Bian, X., and De Camilli, P. (2018). A liquid phase of synapsin and lipid vesicles. Science 361, 604–607.

Mrestani, A., Pauli, M., Kollmannsberger, P., Repp, F., Kittel, R.J., Eilers, J., Doose, S., Sauer, M., Sirén, A.L., Heckmann, M., et al. (2021). Active zone compaction correlates with presynaptic homeostatic potentiation. Cell Rep 37, 109770.

Müller, M., Liu, K.S., Sigrist, S.J., and Davis, G.W. (2012). RIM controls homeostatic plasticity through modulation of the readily-releasable vesicle pool. The Journal of neuroscience : the official journal of the Society for Neuroscience 32, 16574–16585.

Neher, E., and Brose, N. (2018). Dynamically Primed Synaptic Vesicle States: Key to Understand Synaptic Short-Term Plasticity. Neuron 100, 1283–1291.

Nicoll, R.A., and Schmitz, D. (2005). Synaptic plasticity at hippocampal mossy fibre synapses. Nat Rev Neurosci 6, 863–876.

Paul, M.M., Dannhauser, S., Morris, L., Mrestani, A., Hubsch, M., Gehring, J., Hatzopoulos, G.N., Pauli, M., Auger, G.M., Bornschein, G., et al. (2022). The human cognition-enhancing CORD7 mutation increases active zone number and synaptic release. Brain.

Pauli, M., Paul, M.M., Proppert, S., Mrestani, A., Sharifi, M., Repp, F., Kurzinger, L., Kollmannsberger, P., Sauer, M., Heckmann, M., et al. (2021). Targeted volumetric single-molecule localization microscopy of defined presynaptic structures in brain sections. Commun Biol 4, 407.

Pedregosa, F., Varoquaux, G., Gramfort, A., Michel, V., Thirion, B., Grisel, O., Blondel, M., Prettenhofer, P., Weiss, R., Dubourg, V., et al. (2011). Scikit-learn: Machine Learning in Python. J Mach Learn Res 12, 2825–2830.

Pulido, C., and Ryan, T.A. (2021). Synaptic vesicle pools are a major hidden resting metabolic burden of nerve terminals. Sci Adv 7, eabi9027.

Reynolds, E.S. (1963). The use of lead citrate at high pH as an electron-opaque stain in electron microscopy. J Cell Biol 17, 208–212.

Rizzoli, S.O., and Betz, W.J. (2004). The structural organization of the readily releasable pool of synaptic vesicles. Science 303, 2037–2039.

Rollenhagen, A., and Lübke, J.H. (2010). The mossy fiber bouton: the “common” or the “unique” synapse? Frontiers in synaptic neuroscience 2, 2.

Rollenhagen, A., Satzler, K., Rodriguez, E.P., Jonas, P., Frotscher, M., and Lübke, J.H. (2007). Structural determinants of transmission at large hippocampal mossy fiber synapses. The Journal of neuroscience : the official journal of the Society for Neuroscience 27, 10434–10444.

Rostaing, P., Real, E., Siksou, L., Lechaire, J.P., Boudier, T., Boeckers, T.M., Gertler, F., Gundelfinger, E.D., Triller, A., and Marty, S. (2006). Analysis of synaptic ultrastructure without fixative using high-pressure freezing and tomography. The European journal of neuroscience 24, 3463–3474.

Rothman, J.E., Krishnakumar, S.S., Grushin, K., and Pincet, F. (2017). Hypothesis -buttressed rings assemble, clamp, and release SNAREpins for synaptic transmission. FEBS Lett 591, 3459–3480.

Savtchenko, L.P., and Rusakov, D.A. (2007). The optimal height of the synaptic cleft. Proc Natl Acad Sci U S A 104, 1823–1828.

Schindelin, J., Arganda-Carreras, I., Frise, E., Kaynig, V., Longair, M., Pietzsch, T., Preibisch, S., Rueden, C., Saalfeld, S., Schmid, B., et al. (2012). Fiji: an open-source platform for biological-image analysis. Nat Methods 9, 676–682.

Schoch, S., Castillo, P.E., Jo, T., Mukherjee, K., Geppert, M., Wang, Y., Schmitz, F., Malenka, R.C., and Sudhof, T.C. (2002). RIM1alpha forms a protein scaffold for regulating neurotransmitter release at the active zone. Nature 415, 321–326.

Schoch, S., Mittelstaedt, T., Kaeser, P.S., Padgett, D., Feldmann, N., Chevaleyre, V., Castillo, P.E., Hammer, R.E., Han, W., Schmitz, F., et al. (2006). Redundant functions of RIM1alpha and RIM2alpha in Ca(2+)-triggered neurotransmitter release. EMBO J 25, 5852–5863.

Siksou, L., Rostaing, P., Lechaire, J.P., Boudier, T., Ohtsuka, T., Fejtova, A., Kao, H.T., Greengard, P., Gundelfinger, E.D., Triller, A., et al. (2007). Three-dimensional architecture of presynaptic terminal cytomatrix. The Journal of neuroscience : the official journal of the Society for Neuroscience 27, 6868–6877.

Stigloher, C., Zhan, H., Zhen, M., Richmond, J., and Bessereau, J.L. (2011). The presynaptic dense projection of the Caenorhabditis elegans cholinergic neuromuscular junction localizes synaptic vesicles at the active zone through SYD-2/liprin and UNC-10/RIM-dependent interactions. The Journal of neuroscience : the official journal of the Society for Neuroscience 31, 4388–4396.

Tang, A.H., Chen, H., Li, T.P., Metzbower, S.R., MacGillavry, H.D., and Blanpied, T.A. (2016). A trans-synaptic nanocolumn aligns neurotransmitter release to receptors. Nature 536, 210–214.

van Rossum, G., and de Boer, J. (1991). Linking a Stub Generator (AIL) to a Prototyping Language (Python). Paper presented at: EurOpen Conference Proceedings (May 20-24, 1991) (Tromso, Norway).

Vandael, D., Borges-Merjane, C., Zhang, X., and Jonas, P. (2020). Short-Term Plasticity at Hippocampal Mossy Fiber Synapses Is Induced by Natural Activity Patterns and Associated with Vesicle Pool Engram Formation. Neuron 107, 1–13.

Vyleta, N.P., Borges-Merjane, C., and Jonas, P. (2016). Plasticity-dependent, full detonation at hippocampal mossy fiber-CA3 pyramidal neuron synapses. eLife 5, e17977.

Vyleta, N.P., and Jonas, P. (2014). Loose coupling between Ca2+ channels and release sensors at a plastic hippocampal synapse. Science 343, 665–670.

Wang, S.S.H., Held, R.G., Wong, M.Y., Liu, C., Karakhanyan, A., and Kaeser, P.S. (2016). Fusion Competent Synaptic Vesicles Persist upon Active Zone Disruption and Loss of Vesicle Docking. Neuron 91, 777–791.

Wang, Y., Liu, X., Biederer, T., and Sudhof, T.C. (2002). A family of RIM-binding proteins regulated by alternative splicing: Implications for the genesis of synaptic active zones. Proc Natl Acad Sci U S A 99, 14464–14469.

Wang, Y., Okamoto, M., Schmitz, F., Hofmann, K., and Sudhof, T.C. (1997). Rim is a putative Rab3 effector in regulating synaptic-vesicle fusion. Nature 388, 593–598.

Watanabe, S., Rost, B.R., Camacho-Pérez, M., Davis, M.W., Söhl-Kielczynski, B., Rosenmund, C., and Jorgensen, E.M. (2013). Ultrafast endocytosis at mouse hippocampal synapses. Nature 504, 242–247.

Weimer, R.M. (2006). Preservation of C. elegans tissue via high-pressure freezing and freeze-substitution for ultrastructural analysis and immunocytochemistry. Methods Mol Biol 351, 203–221.

Wilke, S.A., Antonios, J.K., Bushong, E.A., Badkoobehi, A., Malek, E., Hwang, M., Terada, M., Ellisman, M.H., and Ghosh, A. (2013). Deconstructing complexity: serial block-face electron microscopic analysis of the hippocampal mossy fiber synapse. The Journal of neuroscience : the official journal of the Society for Neuroscience 33, 507–522.

Wu, X., Cai, Q., Shen, Z., Chen, X., Zeng, M., Du, S., and Zhang, M. (2019). RIM and RIM-BP Form Presynaptic Active-Zone-like Condensates via Phase Separation. Mol Cell 73, 971–984 e975.

Zarebidaki, F., Camacho, M., Brockmann, M.M., Trimbuch, T., Herman, M.A., and Rosenmund, C. (2020). Disentangling the roles of RIM and Munc13 in synaptic vesicle localization and neurotransmission. The Journal of neuroscience : the official journal of the Society for Neuroscience 40, 9372–9385.

Zhang, M., and Augustine, G.J. (2021). Synapsins and the Synaptic Vesicle Reserve Pool: Floats or Anchors? Cells 10, 658.

Zhao, S., Studer, D., Chai, X., Graber, W., Brose, N., Nestel, S., Young, C., Rodriguez, E.P., Saetzler, K., and Frotscher, M. (2012). Structural plasticity of hippocampal mossy fiber synapses as revealed by high-pressure freezing. J Comp Neurol 520, 2340–2351.

